# A fijivirus major viroplasm protein shows RNA-stimulated ATPase activity by adopting pentameric and hexameric assemblies of dimers

**DOI:** 10.1101/2022.04.16.488468

**Authors:** Gabriela Llauger, Roberto Melero, Demián Monti, Gabriela Sycz, Cristián Huck-Iriart, María L. Cerutti, Sebastián Klinke, Evelyn Mikkelsen, Ariel Tijman, Rocío Arranz, Victoria Alfonso, Sofía M. Arellano, Fernando A. Goldbaum, Yann G. J. Sterckx, José-María Carazo, Sergio B. Kaufman, Pablo D. Dans, Mariana del Vas, Lisandro H. Otero

## Abstract

The fijivirus *Mal de Río Cuarto virus* (MRCV) causes a devastating maize disease. Its non-structural protein P9-1, which shows ATPase and RNA binding activities, is the major component of the intracellular viroplasm where virus replication takes place. Here, we established that the 24 C-terminal residues (C-arm) of P9-1 are required for the formation of viroplasm-like structures (VLS) *in vivo* and for the protein multimerization *in vitro*. Employing an integrative structural approach, we found that the C-arm is dispensable for P9-1 dimer assembly, but essential for the formation of doughnut-shaped pentamers and hexamers of dimers (decamers and dodecamers). Both assemblies, larger than those reported for other reoviruses, contain disordered loops oriented towards the inner pore of the structures, where RNA binding sites and conditional proteasome-mediated degradation signals (PEST) were predicted. *In vitro* assays demonstrated that ssRNA binding is favored towards P9-1 (do)decamers over the dimeric ΔC-arm version. In addition, although both P9-1 and P9-1ΔC-arm catalyze the hydrolysis of ATP with similar activity values, an RNA-stimulated ATPase activity was only observed in the full-length protein, indicating a C-arm-mediated interaction between the ATP catalytic site and the allosteric RNA binding sites in the (do)decameric assemblies. Computational studies revealed a stronger preference of phosphate moieties to the decamer in the pore and the C-arm regions, suggesting that the allosteric communication between the ATP and RNA binding sites is favored with this protein arrangement. Overall, our work reveals the structural versatility of a major viroplasm protein providing unprecedented insights into fijivirus viroplasm assembly and function and establishes the structural basis for the development of antiviral strategies against the Mal de Río Cuarto crop disease.

## INTRODUCTION

Plant diseases caused by fijiviruses (family *Reoviridae*) severely threaten crop production. The Mal de Río Cuarto virus (MRCV) is a member of the genus *Fijivirus* (1), which causes the most severe and economically important maize viral disease in Argentina (2), one of the biggest producing and exporting nations worldwide (3). The virus is transmitted by delphacid planthoppers in a persistent-propagative manner (4). Other fijiviruses severely affect rice and maize production in Asia and Europe (5).

Members of the *Reoviridae* family replicate and assemble within membrane-less cytoplasmic inclusion bodies called viroplasms or viral factories. These structures are formed at early time points after infection and are composed of viral proteins and RNA as well as several host factors such as α/β tubulin and nuclear ribonucleoproteins (hnRNPs) (6). Recently, it has been shown that liquid-liquid phase separation (LLPS) underpins the formation of viroplasms (7, 8), including those of rotavirus (9). Importantly, the inhibition of the expression of viroplasm proteins impairs viral replication in plants and animal cells, as reviewed in (6, 10).

*Fijivirus* particles have a double-shelled, icosahedral structure of 65-70 nm in diameter and contain 10 double-stranded RNA (dsRNA) genomic segments that code for at least 12 proteins (1). Viroplasms produced by the *Fijivirus* members *Rice black streaked dwarf virus* (RBSDV) and *Southern rice black-streaked dwarf virus* (SRBSDV) present two distinct morphologies: one granular, predominantly composed by the viral non-structural protein P9-1 and other filamentous, predominantly composed by the non-structural protein P5 (11, 12). The non-structural protein P6 is driven to both types of viroplasms by direct protein-protein interactions with P9-1 and P5 (11, 13). Such structures are highly dynamic: viral RNA has been proposed to accumulate in the granular viroplasm, and viral progeny core and complete virus particles accumulate in the more electrodense filamentous viroplasm, as indicated by the presence of structural proteins P8 and P10 (11).

The MRCV genome codes for six structural proteins (P1-P4, P8, and P10) and six non-structural proteins (P5, P6, P7-1, P7-2, P9-1, and P9-2) (14–18). MRCV P9-1 (hereinafter P9-1) localizes in viroplasms of plant and insect hosts (19) and, when expressed alone, self-interacts giving rise to cytoplasmic viroplasm-like structures (VLS) (20–22). In addition, as shown for animal reovirus counterparts, P9-1 binds single-stranded nucleic acids (ssRNA and ssDNA) in a sequence-independent manner and has ATPase activity. Overall, these properties led us to propose that P9-1 is the major viroplasm component (20). Although the precise role of these enzymatic activities remains unknown, the energy derived from ATP hydrolysis may be required for virus replication and packaging. In turn, MRCV P6 is a minor viroplasm component that forms perinuclear inclusions, is driven to VLS as it interacts with P9-1, and self-interacts through a predicted coiled-coil domain (22, 23). Moreover, both P9-1 and P6 contain proteasome-mediated degradation signal (PEST) sequences that are conserved within fijiviruses (22) and can act as conditional proteolytic signals that target proteins for proteasomal degradation (24).

A few viroplasm proteins from animal and plant reoviruses have been structurally characterized. Studies on rotavirus non-structural protein NSP2 have shown that it is functional as a doughnut-shaped octamer with a central pore and prominent diagonal grooves where NSP5 and ssRNA bind (25, 26). In structural proximity to the RNA-binding grooves, each NSP2 monomer has clefts containing an NTPase active site (27). In turn, the crystallographic structure of the N-terminal domain of Bluetongue virus (BTV, *Orbivirus, Reoviridae*) NS2 revealed that this protein homo-multimerizes through extensive monomer-monomer interactions (28) and electron microscopy studies of the full-length version indicated that the oligomers present a ring-like shape (29). Regarding plant reovirus, cryo-electron microscopy (cryo-EM) analysis of Pns9 from *Rice gall dwarf virus* (RGDV, *Phytoreovirus, Reoviridae*) revealed the formation of octamers with an internal pore (30). Similarly, a crystallographic analysis showed that RBSDV P9-1 forms dimers that interact with each other through carboxy-terminal regions of 24 residues (C-arms) giving rise to cylindrical octamers (31). Residues 25 to 44 are important for RNA binding and reside within a disordered region located at the inner pore (32). Deletion of the C-arm impairs the formation of VLS when expressed in Sf9 insect cells (31). A crystal structure of the SRBSDV P9-1 octamer is available (33), and deletion of the C-arm also affects octamer formation (34). In agreement, we have previously shown that MRCV P9-1 residues 155 to 337 are required for VLS formation in insect cells (20) and deletion of the C-arm affects its self-interactions in yeast two-hybrid (Y2H) assays (22).

The structure and function of viroplasm components underpin the precise coordination of virus replication and packaging. These steps are particularly complex in the case of viruses with segmented dsRNA genomes that require equimolar packaging of all segments. The mechanisms underlying this process are being increasingly understood in animal reoviruses where phosphorylation cascades on rotavirus NSP2 and NSP5 and the RNA chaperone function of NSP2 have crucial roles, as recently reviewed (6, 35). However, far less is known about structure-function aspects of plant reovirus viroplasms.

To shed light on MRCV viroplasm assembly and function, we carried out an integrative structural characterization of P9-1 that evidences that the C-arm-driven oligomerization leads to quaternary structural conformations with an RNA boosted ATPase activity previously unidentified for a major viroplasm protein of the *Reoviridae* family.

## RESULTS

### The P9-1 C-arm is required for the formation of VLS in rice protoplasts and insect cells

We have previously shown that P9-1 forms VLS in the cytoplasm of plant and insect cells (20–22). To assess the contribution of the P9-1 C-arm in the formation of such structures, we transiently expressed P9-1 (337 residues) and P9-1ΔC-arm (lacking residues 314 to 337) fused to the green fluorescent protein (GFP), and analyzed their subcellular localization in rice protoplasts and insect Sf9 cells by confocal imaging. As expected, GFP:P9-1 fluorescence was located in punctuate, distinct cytoplasmic inclusion bodies both in plant and insect cells (**Fig. 1**). Deletion of the C-arm resulted in a dispersed cytoplasmic GFP fluorescence in both systems (**Fig. 1**), indicating that VLS formation was impaired. These results suggest that the C-arm plays a key role during P9-1 multimerization, which is required for VLS formation.

**Figure 1.**
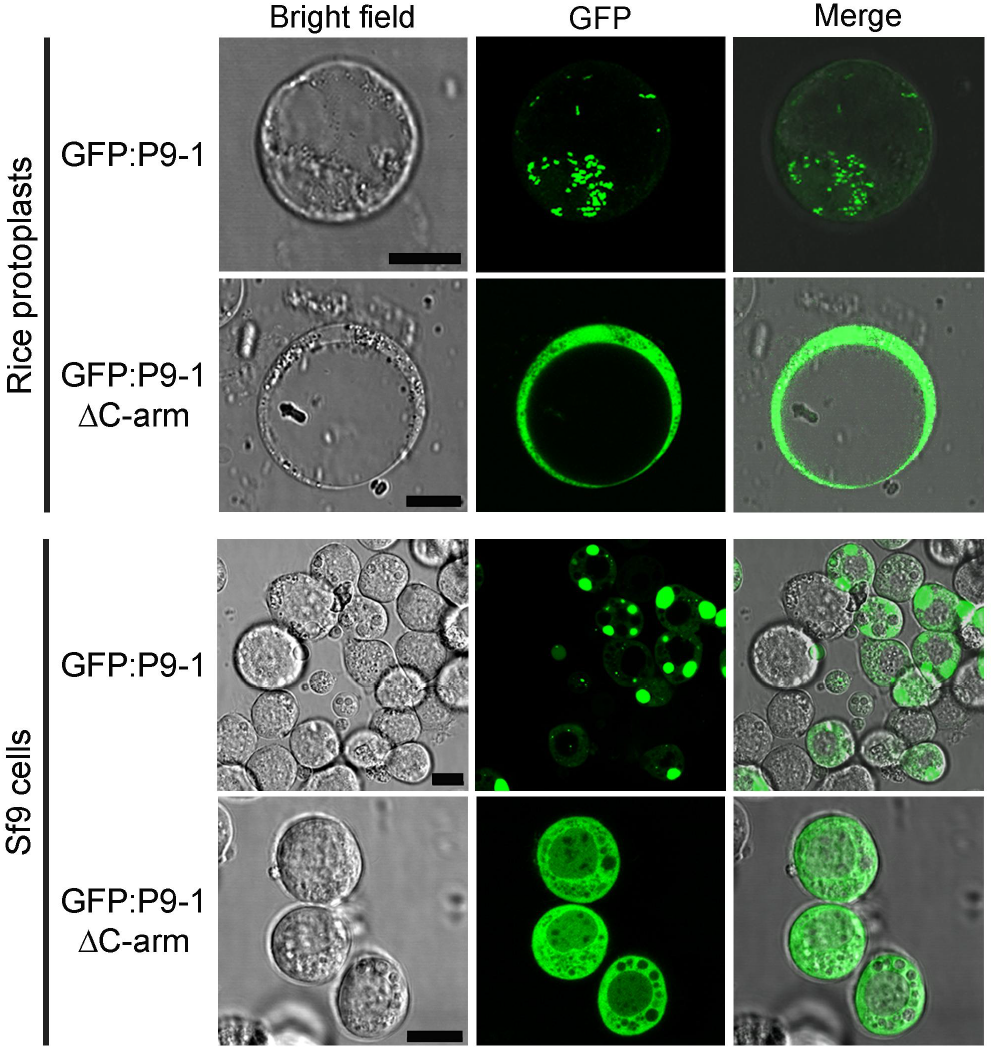
The formation of VLSs in plant and insect cells depends on the presence of the P9-1 C-arm. Confocal microscopy imaging of rice protoplasts (*upper*) or Sf9 insect cells (*bottom*) expressing GFP:P9-1 or GFP:P9-1ΔC-arm. Images of the bright field, GFP fluorescence, and their merge are shown. Scale bars correspond to 10 µm (black bars on bright-field images).

### P9-1 multimerizes into high molecular mass complexes that rely on the presence of the C-arm

To assess whether VLS formation is a result of P9-1 self-interactions leading to multimerization, we subsequently analyzed P9-1 and P9-1ΔC-arm oligomeric states in solution by size-exclusion chromatography (SEC) coupled to static-light scattering (SLS).

Both proteins were produced recombinantly in bacteria and purified by immobilized metal affinity chromatography (IMAC) followed by SEC, and their purity was assessed by SDS-PAGE (**Fig. 2A**). Under reducing conditions, the proteins migrated according to the theoretical molecular mass (MM) of their monomeric species (full-length P9-1, 44.9 kDa and P9-1ΔC-arm, 37.2 kDa). The SEC-SLS analyses showed that P9-1 and P9-1ΔC-arm mostly eluted as oligomeric structures harboring ∼10.4 protomers (experimental MM = 467.9 ± 32.7 kDa) (**Fig. 2B**), and ∼2.3 protomers (experimental MM = 85.3 ± 5.8 kDa) (**Fig. 2C**), respectively. The P9-1ΔC-arm sample was characterized by a persistent high non-specific SLS signal (with low refraction index), which could be indicative of the spontaneous formation of multiple (and subvisible) aggregates possibly due to the instability of the protein construct. Overall, these results indicated that P9-1 behaves as a higher-order oligomer, while the P9-1ΔC-arm variant does not multimerize beyond a dimeric state. These findings are in agreement with those previously reported for RBSDV P9-1, where the C-arm is required for octamer formation but not for the dimer assembly (31).

**Figure 2.**
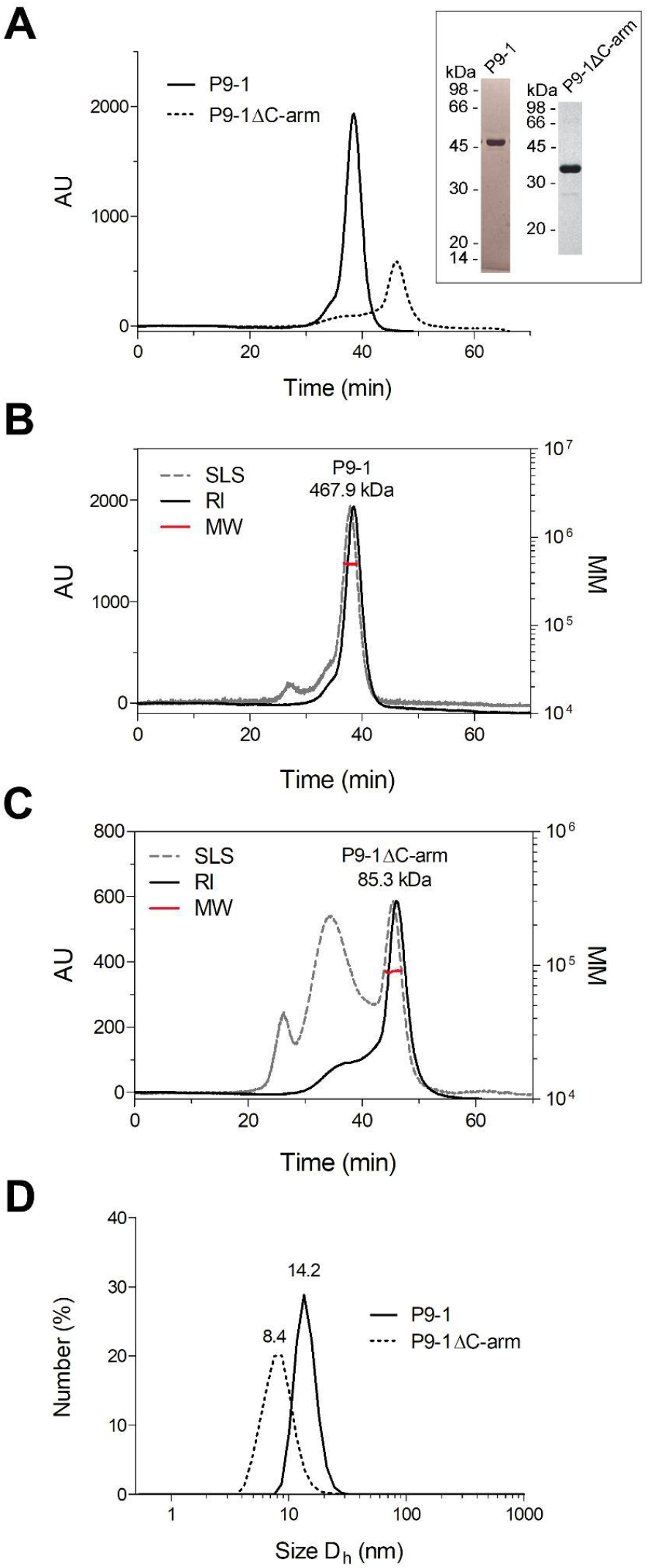
Analyses of P9-1 and P9-1ΔC-arm in solution reveal that the C-arm is required for the assembly of high MM multimers but not for dimerization. **A**. Overlapping SEC chromatograms of P9-1 and P9-1ΔC-arm. SDS-PAGE of the fractions corresponding to P9-1 and P9-1ΔC-arm peaks are shown in the inset. **B** and **C**. SEC-SLS analyses of P9-1 and P9-1ΔC-arm. Normalized light scattering at 90° (SLS, grey dotted line) and refraction index (RI, black line) signals of the eluted proteins for P9-1 (**B**) and P9-1ΔC-arm (**C**) are shown. The trace of the calculated MM is presented in red. A representative run from two independent experiments is shown for each protein, and the average experimental MM value for each protein is indicated above the peaks. **D**. DLS size distributions by number for P9-1 and P9-1ΔC-arm. The number above the peak corresponds to the estimated D_h_. AU, arbitrary units.

To provide further evidence on the oligomerization states of the two proteins, dynamic light scattering (DLS) measurements were performed. P9-1 showed a major population obtained by number distribution corresponding to a hydrodynamic diameter (D_h_) of 14.17 ± 0.87 nm (**Fig. 2D** and **Table S1**), which is consistent with a large multimeric structure, as reported by the SEC-SLS results. No signals compatible with monomeric or low oligomeric states were observed. In turn, the D_h_ obtained by number distribution for P9-1ΔC-arm was 8.41 ± 1.40 nm (**Fig. 2D** and **Table S1**) indicating that the size distribution of the variant assembly was significantly smaller than the full-length protein. In both cases, the percentage of aggregated material present in the analyzed samples was negligible. Complementary analytical SEC studies of P9-1 showed an experimental MM of ∼421.1 kDa (**Figs. S1A** and **S1B**), which is in very good agreement with the estimations based on SEC-SLS experiments. The D_h_ obtained via analytical SEC for P9-1 (∼14.6 nm) is also fully coherent with the value determined via DLS results (**Fig. S1C**). Taken together, SEC-SLS, DLS, and analytical SEC consistently indicated that P9-1 forms higher-order oligomers with stoichiometries that likely exceed an octameric assembly (theoretical MM of 359.2 kDa) as reported for RBSDV P9-1 (31).

### The crystal structure of P9-1ΔC-arm reveals a dimeric arrangement

Attempts to crystallize the full-length P9-1 protein were unsuccessful since they consistently rendered low-quality crystals. Instead, P9-1ΔC-arm crystallized in the tetragonal space group *P*4_3_2_1_2 with unit-cell parameters *a*=*b*= 86.56Å, *c*= 95.60Å, and the best diffraction data set was collected to a maximum resolution of 3.47 Å (**Table S2**). The crystal structure was solved by the molecular replacement method using the atomic coordinates of RBSDV P9-1 as search model (PDB code: 3VJJ) where one independent molecule of P9-1ΔC-arm was found in the asymmetric unit.

The final 2*mF*_o_-*DF*_c_ electron density map was consistent, with no chain breaks for most of the protein backbone, except for the initial four N-terminal residues, and the regions comprising the residues 20-43, 71-72, 108-110, 131-154, 229-237, and 265-268, which correspond mainly to loops. Despite the moderate resolution reached, the final refined model showed good stereochemistry parameters (98^th^ percentile according to MolProbity score (36) on structures of comparable resolution), and acceptable refinement statistics (*R*_work_= 0.22, *R*_free_= 0.28) (**Table S2**).

The P9-1ΔC-arm secondary structure elements comprise nine α-helices and nine β-strands organized as follows: αI (residues 51-61), αII (64-68), αIII (80-101), αIV (173-201), αV (205-209), αVI (211-228), αVII (238-247), αVIII (251-260), αIX (270-277), βA (14-17), βB (45-48), βC (114-117), βD (122-125), βE (156-159), βF (279-286), βG (289-297), βH (303-306), and βI (308-311) (**Fig. 3A**). The longest helix αIV crosses the entire protein fold enclosed by the other α-helices forming a compact helix bundle. Three antiparallel stranded β-sheets constituted by the strands βA(↓),βB(↑) <β-sheet 1>, βC(↑),βD(↓),βE(↑),βI(↓) <β-sheet 2>, and βF(↓),βG(↑),βH(↓) <β-sheet 3> are exposed to the solvent flanking a side of the helix bundle almost perpendicularly with respect to the helix αIV (**Fig. 3A**). The strands βF and βG from β-sheet 3 protrude out from the global protein scaffold as a β-hairpin, while the loops βA-βB (18-44) and βD-βE (126-155) are partially defined by the electron density map revealing local flexibility (**Fig. 3A**). These observations are in line with the prediction of intrinsically disordered regions (IDRs) based on the P9-1 amino acid sequence (**Fig. S2**). Remarkably, the loop βA-βB comprises the RNA binding site previously described for RBSDV P9-1 (residues 25-44), while the loop βD-βE contains the PEST motif (KTESTSSELPAK, residues 142-153) for putative proteasome-mediated degradation (**Fig. S3**).

**Figure 3.**
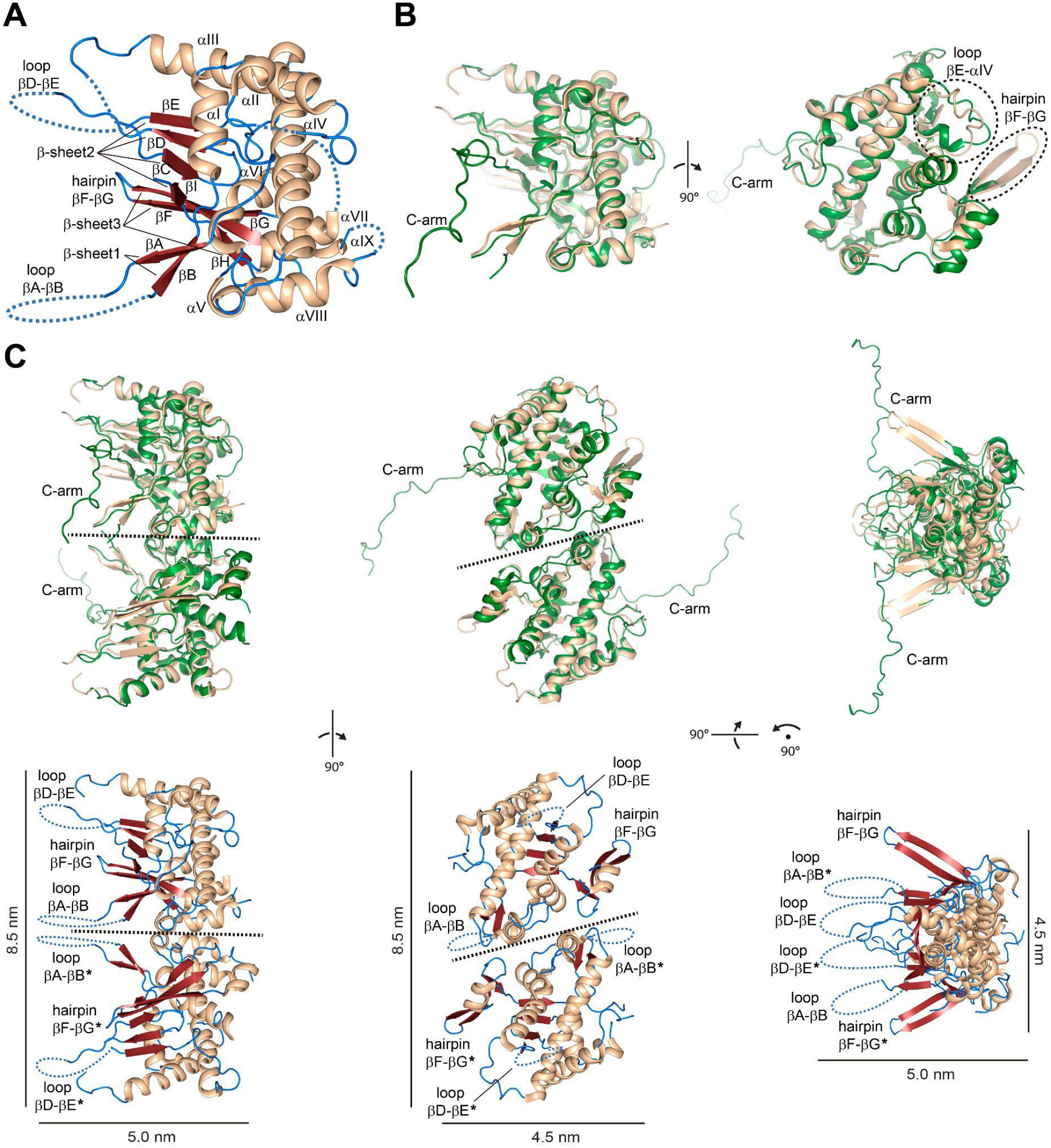
The crystal structure of P9-1ΔC-arm confirms that the C-arm is not crucial for the protein dimer assembly. **A**. Monomer as defined in the asymmetric unit. The secondary structure elements are labeled and colored by type: α-helix, wheat; β-strand, red; loops, blue. **B**. Structural contrast between P9-1ΔC-arm (wheat) and the full-length crystal structure of RBSDV P9-1 (green; PDB code: 3VJJ, chain A) in two orientations. The most prominent differences, which include the protruding hairpin βF-βG (not defined in RBSDV P9-1) and the altered scaffold in the loop βE-αIV, are highlighted by dashed ovals. The C-arm region is indicated in RBSDV P9-1. **C**. Superposition between the non-crystallographic homodimer of RBSDV P9-1 and the crystallographic homodimer of P9-1ΔC-arm constructed by means of the *y, x, -z* symmetry operation of the *P*4_3_2_1_2 space group (*upper*). The structures are colored according to **B**. Dimeric structural assembly of P9-1ΔC-arm colored by the secondary structure elements as in **A** is shown (*bottom*). The hairpins βF-βG and the loops βA-βB and βD-βE from chains A and B (indicated by asterisks), which protrude from the main body of the dimer to nearly the same direction, are labeled. Three different orientations are shown for clarity purposes and a dashed line is depicted in the dimer interface. Curved dashed lines represent the disordered regions undefined in the electron density map. Scale bars are shown.

P9-1ΔC-arm folds nearly similar to RBSDV P9-1 (31), as revealed by an r.m.s.d. of 1.21 Å for 221 aligned C^α^ atoms (**Fig. 3B**). However, some appreciable differences are noted. The protruding hairpin βF-βG is not defined in RBSDV P9-1, while the loop βE-αIV (160-172) in the P9-1ΔC-arm structure shows an altered scaffold mainly due to the absence of an α-turn (**Fig. 3B**).

The RBSDV P9-1 crystal structure revealed two molecules in the asymmetric unit, which form a non-crystallographic homodimer (31). In P9-1ΔC-arm, an identical dimeric arrangement is observed between two protomers belonging to neighboring asymmetric units (**Fig. 3C, *upper***). These polypeptide chains are related by a two-fold symmetry axis and the dimer can be constructed by means of the *y, x, -z* symmetry element of the *P*4_3_2_1_2 space group.

The P9-1ΔC-arm dimeric arrangement, supported by the SEC-SLS and DLS experiments described above (**Figs. 2C** and **2D**), is stabilized by an interface area of 693 Å^2^ (4.9% of the total solvent‐accessible surface per protomer) with a solvation free energy and a total binding energy upon formation of the dimer interface of -4.44 and -9.77 kcal/mol, respectively, which denote protein affinity according to the PDBePISA server (37). In the dimeric assembly, the hairpins βF-βG and the loops βA-βB and βD-βE from both protomers protrude from the main body of the dimer in nearly the same direction (**Fig. 3C, *bottom***). The P9-1ΔC-arm dimer shows a length of 8.5 nm in its largest dimension (**Fig. 3C, *bottom***), which is consistent with the D_h_ of ∼ 8.4 nm estimated by the DLS measurements (**Fig. 2D** and **Table S1**).

The dimer involves 20 interfacing residues per protomer (8.2% of the protein total residues) encompassed in the helices αV, αVII, αVIII, and the loops αVII-αVIII and βG-βH. The residues Arg207 (αV), Asp253, Gln258 (αVIII), Asn249, Tyr250 (loop αVII-αVIII), and Ser302 (loop βG-βH) interact by means of hydrogen bonds (**Fig. 4, *upper left***), while Phe247 (αVII), Pro251 (loop αVII-αVIII), Leu254, Phe257 (αVIII), and Ile299 (loop βG-βH) form the hydrophobic contacts (**Fig. 4, *bottom left***). Most of the atomic contacts across the dimer interface are identical to those observed in full-length RBSDV P9-1 (**Fig. 4, *right***), confirming that this assembly is not impaired by the deletion of the C-arm region.

**Figure 4.**
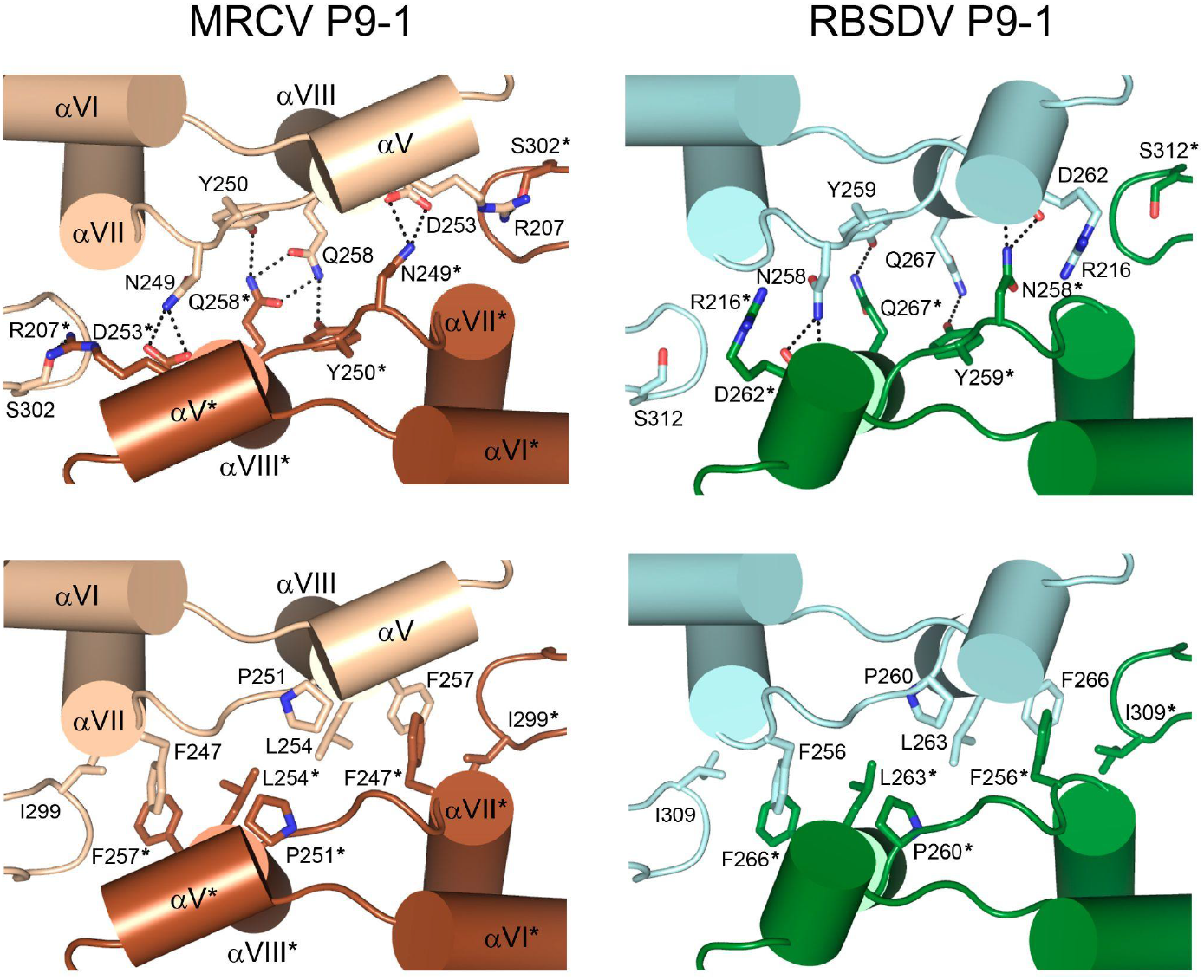
The P9-1ΔC-arm dimer interface is nearly identical to that observed in full-length RBSDV P9-1. Detailed view of the dimerization interfaces from P9-1 (*left*) and RBSDV P9-1 (*right*). Structures are shown in ribbon representation and colored following **Fig. 3B** with the two protomers A and B (indicated by asterisks) depicted in different shades. The most relevant interfacing residues are depicted as sticks and colored according to their corresponding chain. Polar (*upper*) and hydrophobic (*bottom*) interactions are shown. The secondary structure elements from P9-1ΔC-arm involved in the dimer interface are labeled.

### Cryo-EM analysis shows that P9-1 multimerizes as pentamers and hexamers of dimers with an internal pore

Given (i) our previous results in solution in which more complex oligomeric structures were evidenced in the full-length protein (**Figs. 2** and **S1**) and (ii) the fact that RBSDV P9-1 forms an octamer where adjacent dimers are related by a four-fold axis through their C-arms (31), we further performed single-particle cryo-EM studies on the full-length P9-1 protein.

The careful analysis of the recorded data clearly exposed two doughnut-shaped assemblies (torus topology) with ten equivalent subunits (10-mer D5 symmetry) and twelve equivalent subunits (12-mer D6 symmetry), representing 80-85% and 15-20% of the particle populations, respectively (**Fig. 5A**). These data are consistent with the oligomerization state of ∼10.4 protomers estimated by the SEC-SLS assays shown above (**Fig. 2C**).

**Figure 5.**
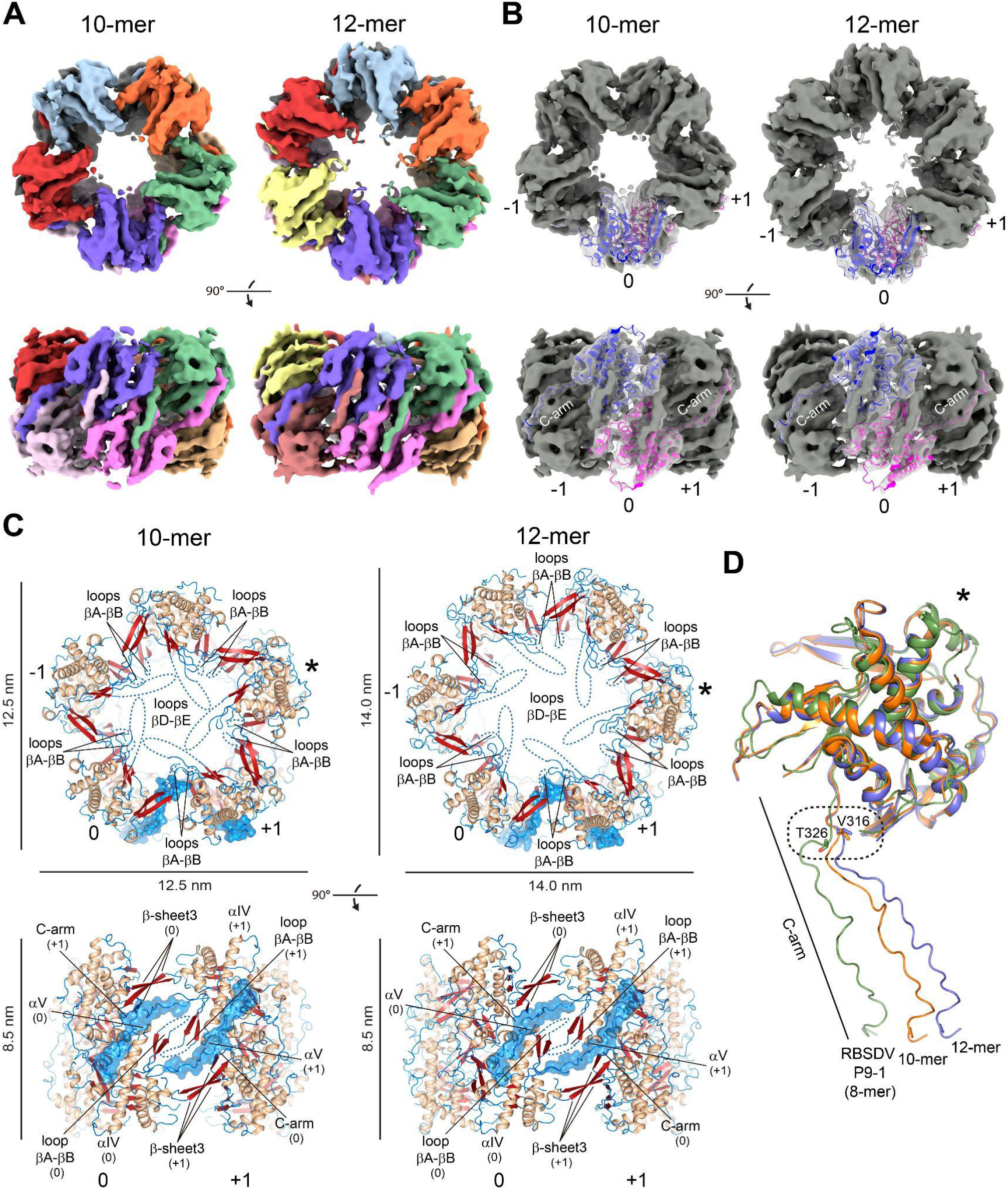
Cryo-EM structures of full-length P9-1 show decameric and dodecameric quaternary arrangements with an internal pore. **A**. Cryo-EM density maps rendered as a surface of the decamer (10-mer) D5 symmetry (*left*) and dodecamer (12-mer) D6 symmetry (*right*) with each structural protomer color-code. Two orientations (top view and side view) of each map or atomic model are shown in **A, B**, and **C. B**. Atomic model of a single full-length dimer (0) fitted in the density maps of the decamer (*left*) and dodecamer (*right*). The model is shown in ribbon representation, colored following the protomer color code of **A**, and displayed with the maps (grey) transparently overlayed. The C-arm density regions and the respective traced chains are indicated. The adjacent dimers are labeled as (−1) and (+1). **C**. Atomic models of the decamer (*left*) and dodecamer (*right*) built from the density maps. The structures are colored according to secondary structure elements as in **Fig. 3A**. The loops βA-βB and βD-βE, which protrude to the internal pore at the middle and the extremes of the structure, respectively, are labeled. Curved dashed lines indicate the disordered regions in the reconstruction of the loops. For clarity purposes, the C-arms are highlighted by the solvent-accessible surface (transparent blue) in the background calculated by PyMOL. Scale bars are shown. Protomers marked with asterisks in **C** and **D** represent analogous views for clarity. **D**. Structural comparison among individual full-length protomers of P9-1 decamer (10-mer, orange), P9-1 dodecamer (12-mer, blue), and RBSDV P9-1 (8-mer, green). A bar indicates the C-arm regions. The change noted in the C-arm trajectory (hinge point) amongst the three promoters is highlighted by a dashed rounded rectangle. The residues found at the dislocation (Val316 in P9-1 and Thr326 in RBSDV P9-1) are depicted as sticks and colored according to their corresponding promoters.

The global resolutions for each species were 4.7 Å (decamer) and 6.8 Å (dodecamer) based on the “gold standard” criterion (FSC=0.143), with local resolutions ranging from 2.5 to 4.5 Å (decamer) and 5.5 to 9.5 Å (dodecamer). We refer the reader to the Materials and Methods section and **Table S3** and **Fig. S4** for details on data acquisition, data processing, and map statistics.

The P9-1ΔC-arm dimer crystallographic structure perfectly fitted as a rigid body into the respective EM-density maps revealing a pentamer of homodimers (decamer) and a hexamer of homodimers (dodecamer) (**Fig. 5B**). In addition, density protrusions corresponding to the C-arm regions (residues 314-337) were clearly distinguishable amongst the docked dimers on both quaternary assemblies (**Figs. 5A** and **5B**). Thus, the respective C-arms were traced and real-space refined in both density maps along with the docked dimer crystallographic structures to obtain the complete atomic models (**Figs. 5B** and **5C**), which showed very good refinement statistics and stereochemistry (**Table S3**).

The doughnut-shaped P9-1 structures show dimensions of 12.5 nm by 12.5 nm by 8.5 nm and 14.0 nm by 14.0 nm by 8.5 nm in the decameric and dodecameric assemblies, respectively (**Fig. 5C**). These structural features agreed with the D_h_ value of ∼14.0 nm estimated by the DLS (**Fig. 2D**) and analytical SEC (**Fig. S1C**).

The full-length dimers are settled in a parallel mode related by the respective five-fold and six-fold rotational symmetry axes holding their C-arm protrusions as staplers, as previously reported for the octameric arrangement of the RBSDV P9-1 crystal structure (31). In this way, the C-arm of the subunit A interacts with the neighboring (−1) dimer, while the C-arm of the subunit B interacts with the other-side (+1) dimer (**Fig. 5B**). The loops βA-βB (RNA binding site) and βD-βE (PEST site), partially defined in the density maps as observed in the P9-1ΔC-arm crystallographic structure, protrude to the internal pore at the central part and the extremes (top and bottom) of the two oligomeric structures, respectively (**Fig. 5C**).

The nascent portion of the C-arm is sandwiched between the β-sheet 3 (hairpin βF-βG + β-strand H) from one adjacent subunit and the loop βA-βB along with the helix αV from the other one, while the distal portion is partially embedded on the surface of the latter nearly aligned to the helix αIV (**Fig. 5C**). According to the PISA server (37), 17 residues (∼70% of the C-arm total extension) are part of the intermolecular contacts with the adjoining dimer, mostly stabilized by hydrophobic forces.

Structural comparison with the RBSDV P9-1 crystal structure revealed a dislocation of the C-arm trajectory with respect to both quaternary assemblies, where the hinge point is noticeable in the nascent C-arm backbone (**Fig. 5D**). Interestingly, despite the high similarity found in the C-arm sequence across homologous proteins, differences are noted in this particular region, where a valine residue (Val316) is found in P9-1 while a threonine residue (Thr326 in RBSDV P9-1) is highly conserved in other *Fijivirus* proteins (**Fig. S3**).

### Small-angle X-ray scattering (SAXS) analysis provides further evidence of the decameric and dodecameric states of P9-1

SAXS was employed to gain further insights into the solution behavior of the P9-1 quaternary assemblies. A thorough analysis of the collected data revealed a small fraction contribution of larger aggregates to the final scattering curve, which was unexpected since no concentration dependency was observed within the concentration range measured. Fortunately, the fraction of larger aggregates was found to be minor, which still allowed us to obtain useful insights.

First, the particle radius of gyration (R_g_) was estimated via Guinier peak analysis (38) to enable the investigation of its compactness and oligomeric state via dimensionless Kratky (39) and power-law (40) analysis, respectively. The R_g_ value obtained from the experimental data (**Fig. 6A**) was estimated at 61.6 Å, which is larger but reasonably close to the R_g_ values obtained from simulated scattering data of the decamer and dodecamer observed in cryo-EM (51.5 and 58.2 Å, respectively) (**Fig. 6B**). The compact, globular nature of the P9-1 higher-order oligomers was confirmed by normalized Kratky analysis (**Fig. 6A, *inset***), and R_g_-based power-law analysis suggests an order of oligomerization of 13, which is fully consistent with the previously described decamer and dodecamer. Second, analysis of the final scattering curve with OLIGOMER (41) revealed that P9-1 decamers and dodecamers were the predominant species in solution (fractions of 87 ± 1% and 13 ± 1%, respectively), with a small contribution by larger aggregates. The decamer proportion decreased to 75 ± 5% when a full pattern modeling was employed (**Fig. 6A**). The two possible oligomers exhibit different features in the Porod region which were useful to estimate the volume fraction of decamers and dodecamers, despite the Guinier region being partially affected by the presence of larger aggregates. The estimation of the MM from the simulated oligomer SAXS patterns (42) using an average density for large proteins (43) was 407 ± 40 kDa for the decamer and 508 ± 50 kDa for the dodecamer, which correspond to values expected for these oligomers, as their theoretical MMs are 449.0 and 538.8 kDa, respectively. Thus, the simulated patterns from cryo-EM density maps were considered representative. The larger aggregates showed a fractal dimension of 2 (platelets-like), which was also observed in other samples with larger aggregates (**Fig. S5A**), where the aggregation degree did not change with dilution (strong particle-particle interaction). There was an additional structural aspect to be considered, as the water-ion affinity may change inside the oligomer pore with respect to the outer protein surface. Thus, solution density inhomogeneity changed the scattering contrast and increased the estimation error of the volume fraction (**Fig. S5B**). Lastly, the final scattering curve was also employed for *ab initio* modeling (**Fig. 6C**). Distance distribution analysis revealed a maximal particle dimension (D_max_) of 183 Å and shape reconstruction with P2 imposed symmetry resulted in a low-resolution model consistent with the dimensions of the decamer and dodecamer observed.

**Figure 6.**
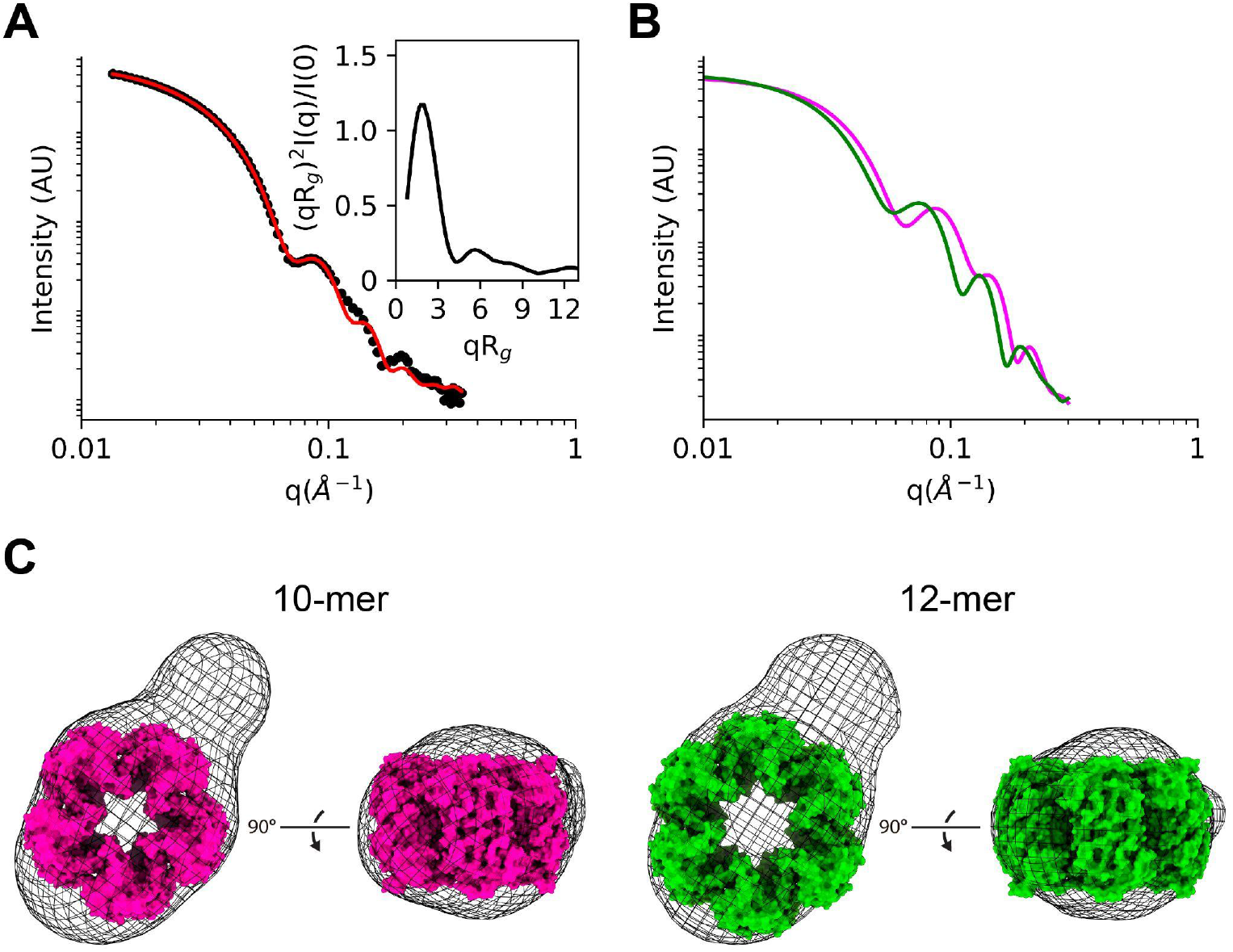
SAXS patterns from cryo-EM density maps corroborate the presence of the decamer and dodecamer assemblies in solution. **A**. Experimental data (dots) and calculated patterns after the non-linear least square procedure. The Kratky plot is shown in the inset. The fitting procedure is detailed in the Materials and Methods section. **B**. Simulated SAXS patterns from cryo-EM map information for decamers (pink line) and dodecamers (green line). AU, arbitrary units. **C**. Low-resolution shape reconstruction yields a particle shape consistent with the dimensions of a decamer (pink) and a dodecamer (green). Two orientations (top view and side view) of each atomic model are shown. The additional unaccounted density results from the above-mentioned minor fraction of larger aggregates that contribute to the scattering curve.

In conclusion, the SAXS data complement the above-mentioned biophysical and structural data and support the simultaneous presence of decameric and dodecameric quaternary P9-1 states in solution, with the former being the predominant species as indicated by cryo-EM.

### P9-1 C-arm-mediated oligomerization into (do)decamers favors RNA binding

To explore whether the nucleic acid binding activity of P9-1 depends on the C-arm, increasing amounts of purified P9-1 and P9-1ΔC-arm were incubated with a 22-mer hexachloro-fluorescein (HEX)-labeled ssDNA probe and subjected to electrophoretic mobility shift assays (EMSA). The migration of the protein-ssDNA complexes was monitored by fluorescence detection of the probe (**Fig. 7A**) and the protein migration by staining the native gel with Coomassie Brilliant Blue (**Fig. 7B**). As expected, P9-1 bound ssDNA in a concentration-dependent manner, showing a statistically significant increment in DNA binding between 3 and 6 µM of protein. Coomassie staining confirmed that the complexes shifted accordingly to the migration of the multimeric assemblies of P9-1. P9-1ΔC-arm also bound ssDNA, and the protein-DNA complexes were less retarded, in agreement with the migration pattern of the dimers formed by this variant protein (**Figs. 7A** and **7B**). Unlike P9-1, ssDNA binding by P9-1ΔC-arm was independent of protein concentration. No statistically significant differences in complex formation beyond 6 µM of protein concentration were observed in either protein (**Fig. 7C**). As a negative control, BSA did not bind ssDNA. These results indicate that the binding of a 22 nt-long ssDNA is independent of the presence of the C-arm (and thus P9-1 higher-order oligomeric states), in line with the potential nucleic acid binding site residing within the loops βA-βB.

**Figure 7.**
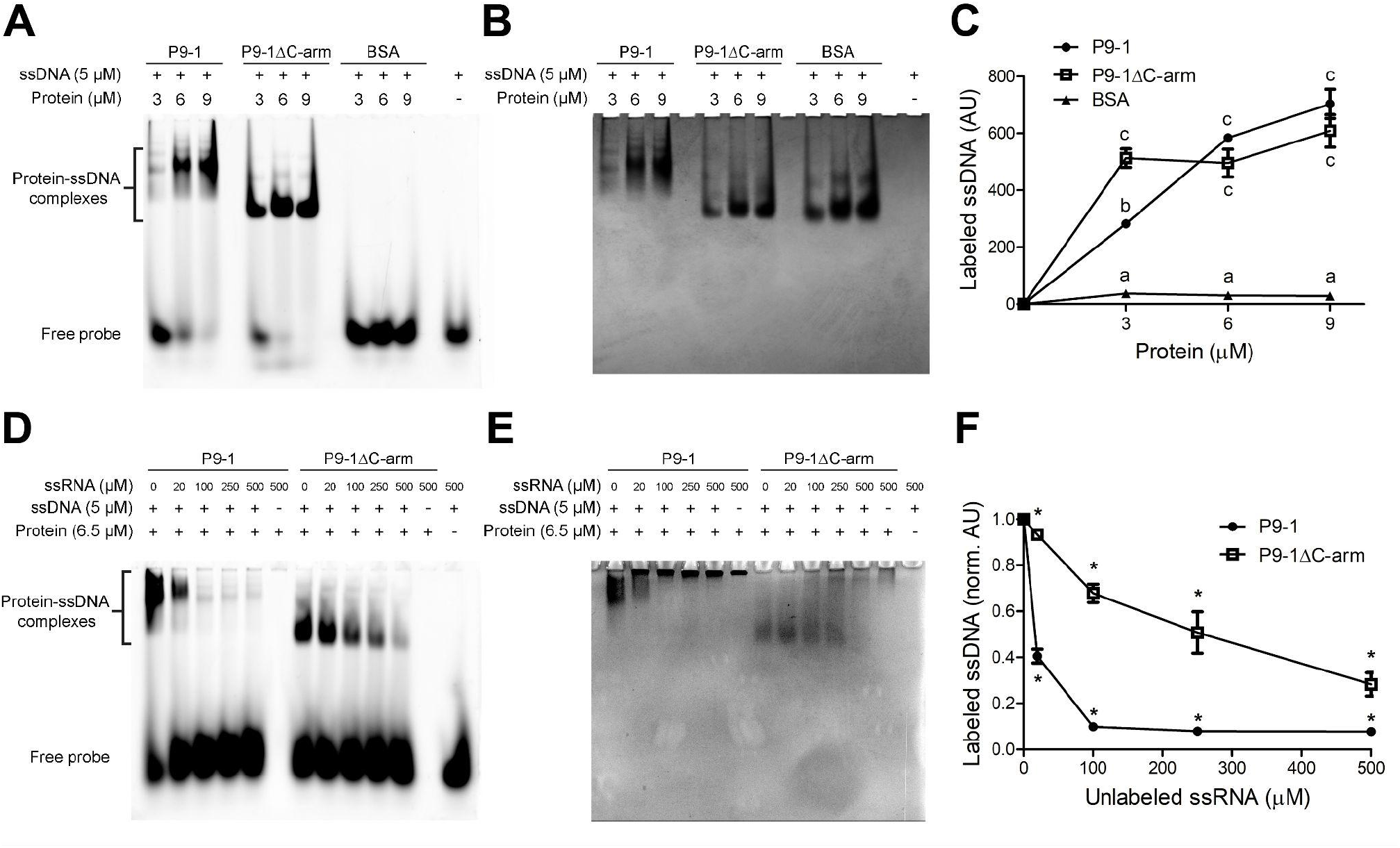
P9-1 preferentially binds ssRNA while deletion of the C-arm does not impair nucleic acid-binding. **(A-C)** EMSA of protein bound to ssDNA. **A**. Native gel of fluorescent ssDNA probe in complex with increasing amounts of P9-1, P9-1ΔC-arm, or BSA, as detected by gel imaging. **B**. The same native gel stained with Coomassie Brilliant blue for protein detection to show the migration patterns of the P9-1 10-mer/12-mer population, P9-1ΔC-arm dimers or BSA (66.5 kDa). **C**. Quantification of the fluorescence intensity of the labeled ssDNA (in arbitrary units, AU) in complex with the proteins. **(D-F)** EMSA of competition assays between labeled ssDNA and unlabeled ssRNA in complex with P9-1 or P9-1ΔC-arm. **D**. Native gel of fluorescent ssDNA probe in the presence of increasing amounts of unlabeled ssRNA, as detected by gel imaging. **E**. The same native gel stained with Coomassie Brilliant blue for protein detection to show the migration patterns of the nucleic acid/protein complexes. **F**. Quantification of the fluorescence intensity of the labeled ssDNA bound to the proteins (in normalized arbitrary units, norm. AU) in the presence of increasing amounts of unlabeled ssRNA. In **C** and **F** each point represents the mean and standard deviation of three independent experiments (n=3). Two-way ANOVA followed by Tukey’s multiple comparison test was performed. In **C**, different letters denote statistically different values, while in **F**, statistically different values between proteins at each ssRNA concentration means are denoted by asterisks (*p*-value < 0.05).

Since ssRNA binding activity is crucial for reovirus replication within the viroplasms, we next performed competition assays by adding increasing amounts of unlabeled ssRNA to the P9-1:ssDNA and P9-1ΔC-arm:ssDNA complexes. Competition EMSAs showed that both proteins can bind ssRNA, but P9-1 binding is more efficient (**Figs. 7D** and **7E**). Semi-quantitative analysis of the labeled ssDNA band patterns revealed a 10-fold decrease in the P9-1:ssDNA complex band at 100 µM ssRNA, as opposed to a lower 1.5-fold decrease for the P9-1ΔC-arm:ssDNA complex (**Fig. 7F**). Interestingly, P9-1 presented a retarded migration pattern at increasing ssRNA concentrations (**Fig. 7E**), suggesting the occurrence of inter (do)decameric interactions.

Overall, our results indicate that the presence of the C-arm strongly favors RNA binding, probably as a result of P9-1 oligomerization into (do)decamers.

### The ATPase activity is stimulated by the binding of RNA to P9-1 (do)decamers

Since it is known that MRCV P9-1 catalyzes ATP hydrolysis (20), we set out to quantitatively determine whether deletion of the C-arm affects the ATPase activity, as well as whether the binding of ssRNA to P9-1 and P9-1ΔC-arm has an effect on such catalytic activity. At a protein concentration similar to the one used in the nucleic acid-binding assays (6.5 μM), both proteins presented similar enzymatic activity values (**Table 1** and **Fig. S6**) whereas non-enzymatic hydrolysis was negligible. Interestingly, at 500 μM ssRNA, the P9-1 ATPase activity increased five times while no detectable effect was observed with P9-1ΔC-arm. These results are indicative of an interaction between the RNA and ATP binding sites that results in an RNA-dependent ATPase activity enhancement in P9-1 (do)decamers that is dependent on the presence of the C-arm.

**Table 1.**
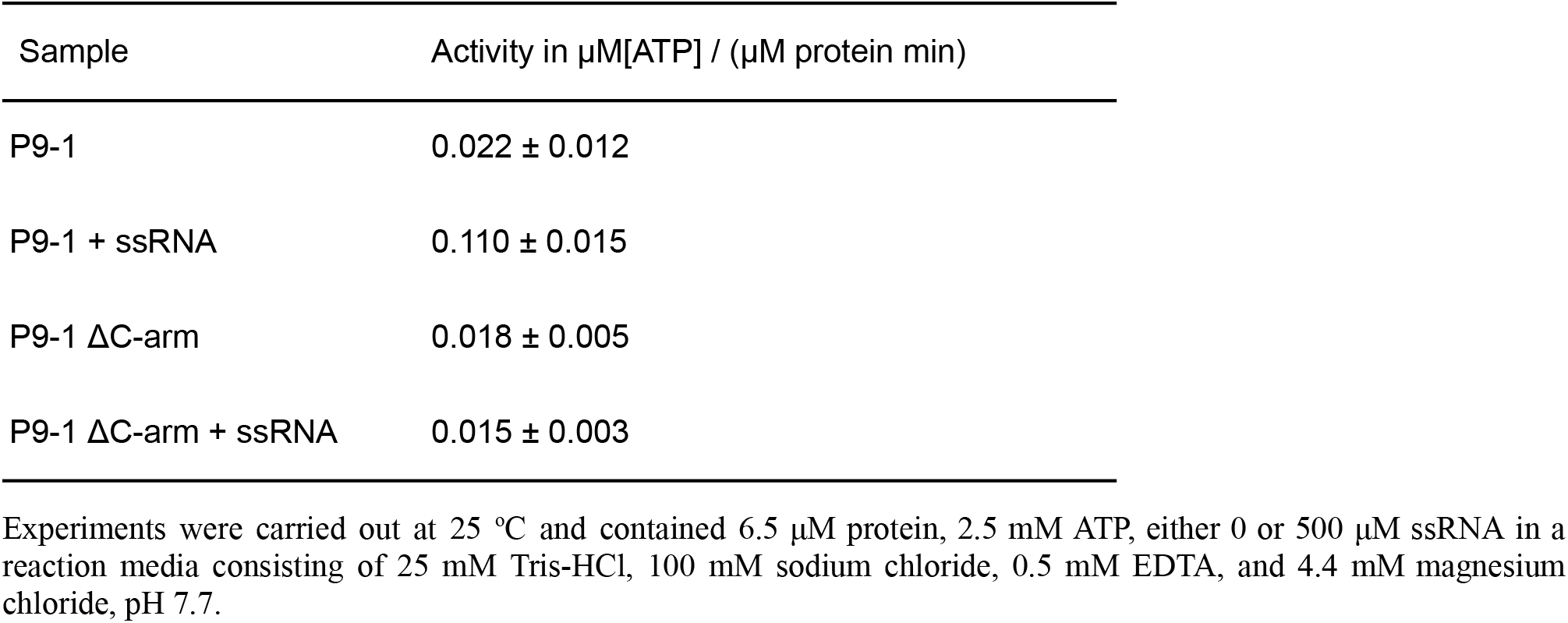
ATPase activity of P9-1 and P9-1ΔC-arm in the presence or absence of ssRNA.

| Sample | Activity in $\mu\text{M}[\text{ATP}] / (\mu\text{M protein min})$ |
| --- | --- |
| P9-1 | $0.022 \pm 0.012$ |
| P9-1 + ssRNA | $0.110 \pm 0.015$ |
| P9-1 ΔC-arm | $0.018 \pm 0.005$ |
| P9-1 ΔC-arm + ssRNA | $0.015 \pm 0.003$ |

Experiments were carried out at 25 °C and contained 6.5 $\mu\text{M}$ protein, 2.5 mM ATP, either 0 or 500 $\mu\text{M}$ ssRNA in a reaction media consisting of 25 mM Tris-HCl, 100 mM sodium chloride, 0.5 mM EDTA, and 4.4 mM magnesium chloride, pH 7.7.

### *In silico* simulations are compatible with a strong binding of phosphate to the P9-1 pore and C-arm that is enhanced in the decameric form

To further characterize the RNA binding and ATPase activities we computed classical molecular interaction potentials on the P9-1 dominant quaternary conformations using spherical probes mimicking the phosphate groups found in RNA and ATP.

First, we rebuilt by means of molecular modeling the missing regions not determined experimentally (see Materials and Methods section) and ran state-of-the-art Molecular Dynamics (MD) simulations in the microsecond timescale. Two dominant 3D conformations of the missing regions (found 35% and 31% of the time for dimers D1; 21% and 19% for decamers D5; and 22% and 21% for dodecamers D6), emerged from the simulations (**Fig. 8A** and **Movies S1-S3**), based on cluster analyses of the positional fluctuation of the molecular systems over time. As above defined, the missing regions are mostly composed of the flexible loops βA-βB and βD-βE that protrude to the internal pore of the (do)decameric assemblies. Interestingly, as a result of the loops orientations, the pore is more occluded in the decamer than in the dodecamer (**Fig. 8A, *middle and right***).

**Figure 8.**
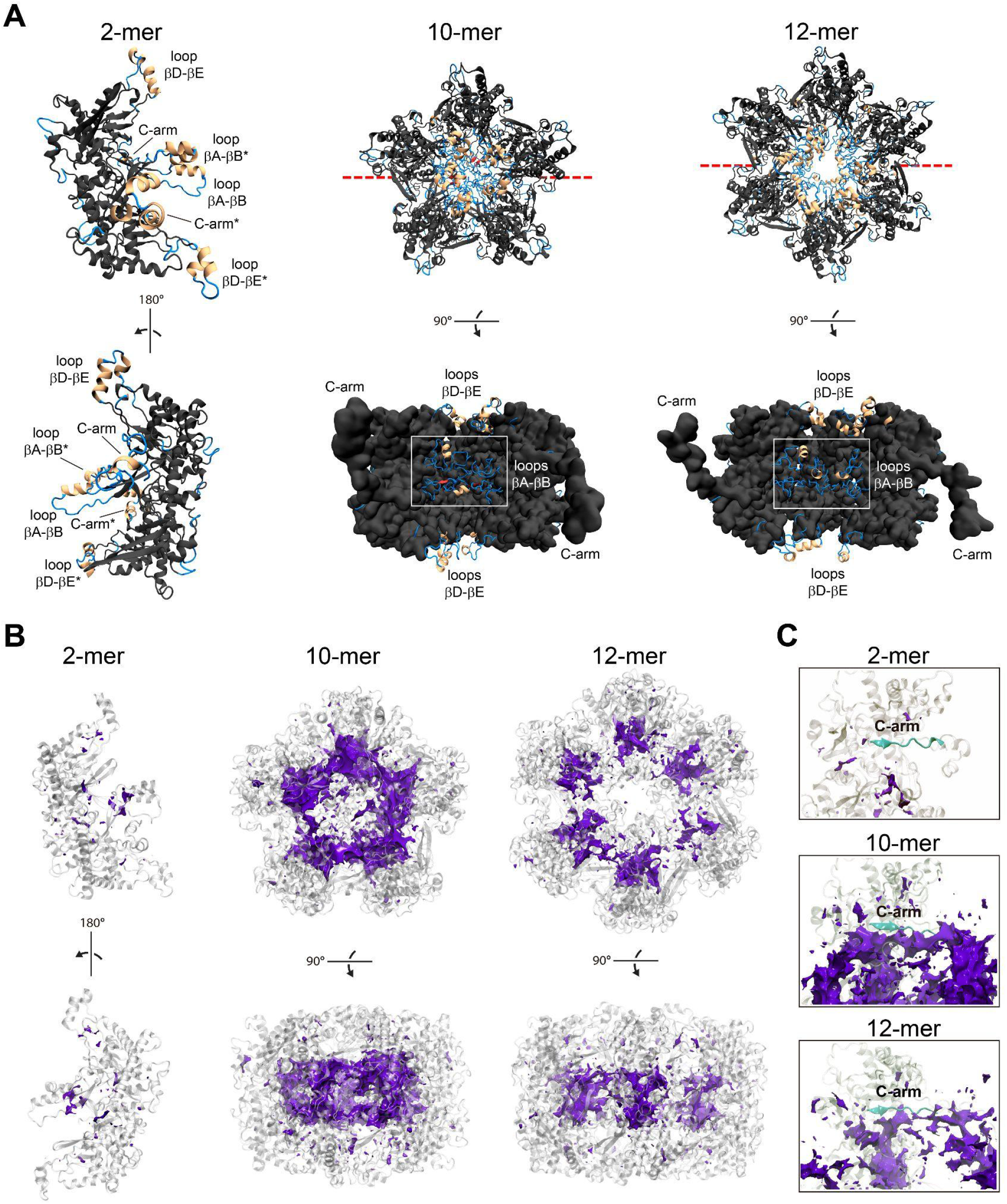
P9-1 flexible loops βA-βB facing the internal pore and the C-arm exhibit marked ability to bind phosphate. **A**. Theoretical reconstructions of P9-1 missing experimental regions in the dimer (*left*), the decamer (*middle*), and the dodecamer (*right*). The representative structure of the dominant conformation extracted after convergence of multi-microsecond MD simulations is shown in each assembly. Rebuilt regions are colored according to secondary structure elements as in **Fig. 3A**; experimental regions are colored in gray. The representation of the dimer assembly is shown in two different orientations. The loops βA-βB and βD-βE and the C-arms from chains A and B (indicated by asterisks) are labeled. The representations of the decamer and the dodecamer are shown in a parallel view to the inner pore (top view) and perpendicular view (bottom view, experimental regions as Connolly surface) with a clipping plane located at the red dashed line. The loops βA-βB and βD-βE (which protrude to the internal pore at the middle and the extremes of both structures, respectively) and the C-arms, are labeled. **B**. Molecular interaction potentials using a phosphate group with charge -1 as a probe in the dimer (*left*), the decamer (*middle*), and the dodecamer (*right*). The isosurface of value -5 kcal/mol is shown with a violet solid mesh in each assembly. Two orientations of each assembly are shown. **C**. Same as **B** but showing the isosurface located on the top and around the C-arm regions (depicted in cyan) in each assembly.

We found a marked ability of P9-1 to bind phosphate moieties in the three multimeric states, with a strong preference for the decamer (**Fig. 8B**). The most probable binding sites found at the -5 kcal/mol isosurface (enthalpic interaction energy) are located inside the pore involving the loops βA-βB, around the new modeled missing regions. Remarkably, a significant binding probability is found on top and around the C-arm region, which is again enhanced in the decameric form (**Fig. 8C**).

## DISCUSSION

Biochemical and structural analyses of reoviral viroplasm proteins are beginning to unravel viroplasm maturation and dynamics. Within the *Reoviridae* family, the major viroplasm proteins from rotavirus (NSP2), phytoreovirus RGDV (Pns9), and fijivirus RBSDV (P9-1) and SRBSDV (P9-1) present ring- or doughnut-shaped octameric structures (25, 30, 31, 33). The results presented in this work show that MRCV P9-1 gives rise to pentamers and hexamers of dimers (10-mers and 12-mers, respectively), which still resemble the overall quaternary structure folding previously reported. Therefore, the functional molecular organization of *Reoviridae* major viroplasm proteins would be conserved, irrespectively if they form octamers, decamers, or dodecamers. Interestingly, the P9-1 C-arm does not affect dimer assembly but instead is critical for the formation of the decameric and dodecameric quaternary structures. Our findings led us to propose a model for the arrangement of the oligomers. Initially, two monomers of P9-1 would interact to form a dimeric assembly across a surface of 20 residues stabilized mainly by hydrogen bonds and hydrophobic interactions. The P9-1 dimer interface, almost identical to that reported in full-length RBSDV P9-1 (31) and SRBSDV P9-1 (33), validates that the dimer assembly is independent of the C-arm and suggests it would be conserved in major viroplasm proteins from plant reoviruses. Next, five or six dimers would interact through their C-arms to give rise to the (do)decameric structures. Both quaternary assemblies are present in an equilibrium where the 10-mers are more abundant than the 12-mers, with no intermediate oligomeric states of lower molecular masses like the dimeric and tetrameric species observed in RBSDV P9-1 (31, 32). We suggest that this versatility in allowing five or six dimers to assemble is given by the flexible nature of the C-arm, possibly caused by a valine residue (Val316), which is only present in MRCV amongst fijivirus. In line with our results, the structural characterization of the full-length BTV NS2 protein by negative stain EM revealed ring-like assemblies that could correspond to decamers or dodecamers (29), suggesting that these higher-order oligomeric structures may also occur in other reovirids.

We have previously reported that the MRCV P9-1 C-terminal half (residues 157-337) is indispensable and sufficient for the formation of VLS (20). The present work allowed us to strengthen this finding by showing that all residues involved in the dimeric and (do)decameric interfaces reside within this region, evidencing its predominant structural role in the protein quaternary assembly.

*Reoviridae* members replicate and package their segmented RNA genome within the viroplasm in a process that is orchestrated by precise intermolecular RNA-RNA interactions and interactions between the viral viroplasm protein components and RNA (35, 44). RBSDV P9-1 preferentially binds ssRNA in the octameric conformation (32) where residues 25 to 44, located in the inner pore of the structure, are necessary for this function. Although residues 319-327 (mostly placed in the C-arm) were also predicted to bind RNA, this could not be entirely assessed since its deletion affects octamer formation (32). Given that P9-1 residues 25-44 and 309-317 align with the RNA binding sites identified for RBSDV P9-1 (**Fig. S3**), we surmise that the P9-1 ssRNA binding activity also resides within these two regions. In P9-1, the residues 25-44 are located within the highly flexible loop βA-βB facing toward the internal pore, while the second predicted RNA binding site (residues 309-317) is located in the nascent C-arm region near the loop βA-βB in the decamer and dodecamer (**Figs. 5C** and **S3**).

As defined by the molecular interaction potentials, both the loops βA-βB and the C-arms contain probable phosphate binding sites, which is indicative of interaction with RNA and/or ATP molecules. Since P9-1ΔC-arm showed similar ATPase activity with respect to the full-length protein, the ATP catalytic site would not reside within the C-arm; therefore, the phosphate moieties detected around this region would exclusively correspond to those from RNA. Moreover, the fact that the absence of the C-arm does not abolish RNA binding but its presence strongly favors it, suggests the existence of RNA binding sites, some of which are dependent (while others are independent) of the C-arm. Altogether, our experimental and theoretical findings are consistent with the previous RBSDV P9-1 observations (32) and support the surmise that the RNA binding site(s) may be located at the loop βA-βB and also at the C-arm. Additionally, because of the proximity of the loops βA-βB and the C-arms within the (do)decameric structures, it is possible to reason that RNA interaction with P9-1 may involve both regions. Future structural studies on P9-1 complexed with RNA will define the specific nucleic acid-binding site(s).

Interestingly, RNA is a non-essential activator of the ATPase activity of P9-1, but the deletion of the C-arm suppressed this activation. Therefore, a structural communication pathway between the ATPase catalytic site and the RNA binding site(s) should exist, which is mediated by the C-arm. This observation suggests that structural changes associated with the interaction of the C-arm with RNA may allosterically modulate the ATPase catalytic site. However, as the C-arm takes up a crucial role in the protein oligomerization into (do)decamers, the disruption of the interaction between both sites observed in P9-1ΔC-arm may be also a consequence of the lack of higher-order oligomers formation. In contrast to the RNA-stimulated ATPase activity observed in P9-1, NSP2 has both NTPase and RNA triphosphatase (RTPase) activities, and RNA competes with NTPs for the active site (27) suggesting they may have different modes of action. Overall, P9-1 ATPase activity, RNA binding, in addition to its quaternary structures resembling the ring-like shape of hexameric helicases (45), support a key role for this protein in MRCV genome replication, reassortment, and packaging.

The different quaternary arrangements would significantly affect the functional communication between the catalytic (ATP) and allosteric (RNA) binding sites, as different phosphate interaction potentials were determined in these regions in the three multimeric states (dimer, decamer, and dodecamer). In this regard, the crowding effect within the pore that takes place in the decamer would benefit the interaction between both sites, as additional and stronger phosphate binding sites were determined. This finding gives a potential functional advantage for the decamer over the dodecamer and the dimer (in that order). The binding preference of phosphate moieties for the decamers may be endorsed by the cryo-EM and SAXS experimental data, which showed that decamers are the most represented assemblies of P9-1 found in solution.

It has been recently shown that LLPS underpins the formation of rotavirus viroplasms (9) and this process is known to be favored by the presence of IDRs and by RNA binding (7, 8). Interestingly, P9-1 contains two IDRs (**Fig. S2**), and the structural models herein described confirm that they correspond to the flexible loops βA-βB and βD-βE. These results, together with the dynamics of the formation of VLS upon P9-1 expression in insect and plant cells (**Fig. 1**), support the hypothesis that P9-1 as well undergoes LLPS for viroplasm formation. Future work will deepen into this important point.

For Rotavirus and avian reovirus, viroplasm formation requires a functional proteasome (46–48). Both P9-1 and the minor viral component of the viroplasm P6 contain PEST motifs proposed to regulate MRCV viroplasm formation and dynamics by controlling protein abundance (22). PEST motifs are stretches of residues rich in proline (P), glutamic acid (E), serine (S), and threonine (T), that act as conditional signals for the ubiquitin 26S-proteasome system (UPS)-mediated degradation and can be activated by different mechanisms such as light, ligand binding or phosphorylation (24). Remarkably, the location of P9-1 PEST motifs within the flexible loops βD-βE, exposed towards the top and bottom of the inner pore of the (do)decameric structures, suggest that changes in phosphorylation may readily affect the conformational stability of the region, accelerating P9-1 UPS-mediated degradation (49). In turn, PEST phosphorylation might be regulated by P9-1 interactions with other proteins such as P6, with RNA or be favored in one of the different P9-1 conformations. Additional research is required to assess the precise mechanism that orchestrates this process.

In conclusion, the work presented here describes for the first time at the molecular level, that a fijivirus major viroplasm component forms doughnut-shaped decamers and dodecamers assembled by pentamers and hexamers of dimers, which are crucial for the potential allosteric communication amongst the catalytic ATP sites and the RNA binding sites. These results illustrate the versatility of the quaternary arrangements of a viroplasm protein and raise the intriguing question of whether these homo-oligomeric structures have different biological properties during the virus infection cycle. Furthermore, our work sets up the structural basis for the development of novel antiviral strategies for the Mal de Río Cuarto maize disease.

## MATERIALS AND METHODS

### Cloning of expression plasmids

The reported pRSET P9-1 construct containing the P9-1 coding sequence (Uniprot accession number D9U542; 337 residues, 39 kDa) in frame with a sequence coding a 52 residue N-terminal tag (6xHis/Xpress/enterokinase cleavage recognition sequence, EK) was employed for P9-1 recombinant expression and purification in bacteria (20). The previously reported P9-1 biochemical characterization including homomultimerization, ATPase, and ssRNA binding activities were performed with this recombinant construct, which codes for a 44.9 kDa protein (20). Next, this construct served as a template for a PCR designed to clone the P9-1ΔC-arm (residues 1 to 313 of P9-1, lacking 24 residues at the C-terminus) with an N-terminal 6xHis tag into the pET24a vector. The previously described pCR8/GW/TOPO (Invitrogen, USA) entry vectors containing the P9-1 and P9-1ΔC-arm (22) coding sequences were used for recombination with the LR Clonase II enzyme mix (Invitrogen, USA) according to the manufacturer’s instructions. For live imaging in transfected rice protoplasts, the pUC57-43 vector was employed (50), whereas, for live imaging in transfected Sf9 insect cells, the pIB-GW destination vector (21) was employed. The resulting constructs express P9-1 or the variant P9-1ΔC-arm (lacking the 24 C-terminal residues), fused to the green fluorescent protein at the N-terminus (GFP: P9-1 or GFP: P9-1ΔC-arm).

### Rice protoplasts and Sf9 cells transfection and fluorescence live imaging

Rice japonica variety Kitaake protoplasts were prepared and transfected as described elsewhere (51). *Spodoptera frugiperda* Sf9 (IPLBSF21-AE clonal isolate 9) cells were cultured and transfected as previously described (23). Fluorescence imaging was performed in a Leica TCS-SP5 (Leica Microsystems GmbH, Germany) spectral laser confocal microscope using a 63x objective (HCX PL APO CS 63.0 1.20 WATER UV). The 488 nm line of the Argon laser was used for GFP excitation, and the fluorescence emission was detected with the channel settings 498–540 nm for GFP.

### Protein production and purification

The P9-1 and P9-1ΔC-arm constructs for bacterial expression were transformed into *E. coli* BL21 cells grown in Terrific Broth culture media supplemented with 0.1% glucose and 50 µg/ml ampicillin and induced for expression with 1 mM isopropyl-β-D-thiogalactopyranoside (IPTG) at 28 °C, overnight. Cells were harvested by centrifugation for 15 min at 5000 ×g and 4 °C and resuspended in lysis buffer (20 mM sodium phosphate, 500 mM sodium chloride, 20 mM imidazole, 1 mM phenylmethylsulfonyl fluoride (PMSF), 0.05% Triton X-100, 100 µg/ml lysozyme, pH 7.4) using 10 ml of lysis buffer per 100 ml of cell culture. Soluble proteins were obtained by sonication with 3 pulses of 30 s each in an ice bath using a Vibra-Cell™ Ultrasonic Liquid Processor (Sonics & Materials, Inc., USA) and centrifuged at 12,000 ×g for 15 min at 4 °C. A second extraction was performed by resuspending the remaining pellet in 5 ml of lysis buffer per 100 ml of cell culture. Protein extract was subjected to IMAC by incubation with 2 ml Ni-NTA resin (Qiagen, Germany) per 50 ml of extract, for 3 h at 4 °C with gentle agitation. After incubation, the resin was loaded on an empty column, washed with lysis buffer, and the bound protein was eluted with 20 mM sodium phosphate, 500 mM sodium chloride, 500 mM imidazole, pH 7.4. Buffer exchange and protein sample concentration were performed using 10 kDa MWCO Vivaspin Turbo centricons (Sartorius, Germany). After IMAC, proteins were further purified by SEC using a Superdex 200 column (GE Healthcare, USA), in running buffer (10 mM Tris-HCl, 25 mM sodium chloride, pH 7.6), at a flow rate of 1.3 ml/min, followed by another concentration step with 10 kDa MWCO Vivaspin Turbo centricons. Protein quantification was assessed using a spectrophotometer (NanoDropTM 1000, Thermo Fisher Scientific, USA).

### SEC - SLS measurements

The average MMs of P9-1 and P9-1ΔC-arm in solution were determined on a Precision Detectors PD2010 90º light scattering instrument tandemly connected to high-performance liquid chromatography and an LKB 2142 differential refractometer. The chromatographic runs were performed in a Superdex 200 GL 10/300 column (GE Healthcare) with a buffer containing 10 mM Tris-HCl, 25 mM sodium chloride, pH 7.6 at a flow rate of 0.4 ml/min. The elution was monitored by measuring its SLS signal at 90° and its refractive index (RI). The mass of the injected samples was 150 μg for P9-1 and 300 μg for P9-1ΔC-arm. The MM of each sample was calculated relating its SLS and RI signals and comparison of this value with the one obtained for bovine serum albumin (MM: 66.5 kDa) as a standard. Data were analyzed with the Discovery32 software supplied by Precision Detectors. The average MM value corresponded to the central 10% of the peak.

### DLS measurements

The size distribution and hydrodynamic diameter measurements were performed at 25 ºC with a Zetasizer Nano-S DLS apparatus (Malvern Instruments Ltd., UK) using a low volume quartz cuvette. Protein samples were diluted to ∼2 mg/ml in 25 mM Tris-HCl, 100 mM sodium chloride, pH 7.5. For each sample, 7 to 10 runs of 10 s were performed. Size distributions and hydrodynamic diameters were calculated using the multiple narrow distribution analysis models of the DTS v.7.11 software (Malvern Instruments Ltd., UK).

### Analytical SEC

Analytical SEC was performed using an ENrich 650 10/30 column (Bio-Rad, USA) pre-equilibrated in running buffer (25 mM Tris-HCl, 100 mM sodium chloride, pH 8.0). Bio-Rad gel filtration calibration standard comprising bovine thyroglobulin (670 kDa), bovine γ-globulin (158 kDa), chicken ovalbumin (44 kDa), horse myoglobin (17 kDa), and vitamin B12 (1.35 kDa) was used as MM standard, although the elution volume of the latter was excluded from the analysis. The P9-1 protein sample (500 µl) was injected at 1 mg/ml and eluted at a flow rate of 0.5 ml/min. Calibration of the column was performed using the Bio-Rad MM standard under the same conditions, and the apparent MM of P9-1 was determined according to (52). The partition coefficient (K) was calculated as *K = (V*_*x*_ *- V*_*0*_*)/(V*_*e*_ *- V*_*0*_), where V_x_ is the elution volume of each standard protein, V_0_ is the void volume and V_e_ is the end volume of the column. Estimation of the experimental D_h_ of P9-1 was based on the elution volumes and the D_h_ of the standard proteins, given by the relationship *1000/Vx = a × D*_*h*_ *+ b*.

### Crystallization, X-ray data collection, and structure resolution of P9-1ΔC-arm

Initial crystallization conditions for P9-1ΔC-arm were screened at room temperature on 96-well sitting-drop Greiner 609120 plates using a Digilab Honeybee963 robot (Marlborough, USA) and commercial kits from Jena Bioscience (Jena, Germany) and Hampton Research (Aliso Viejo, USA) at 15 mg/ml protein concentration. Optimized crystals were then grown in 24-well hanging-drop Hampton Research VDX plates with a precipitation solution consisting of 13% (w/v) PEG 8000 and 0.2 M calcium acetate, reaching a maximum size of 0.4 × 0.1 × 0.1 mm^3^. Several detergents and additives (Hampton Research) were tested around this condition, but none of them significantly improved the crystal size and/or diffraction quality. Crystals were cryoprotected in mother liquor supplemented with 22% (w/v) PEG 400 and then flash-cooled in liquid nitrogen using Hampton Research loops.

X-ray diffraction datasets were collected at 100 K on several crystals at the PROXIMA-2A beamline at Synchrotron SOLEIL (France) using an EIGER X 9M detector (Dectris, Baden, Switzerland) and the MXCuBE application (53). The best crystal diffracted to 3.47 Å resolution (**Table S2**). Datasets were indexed, integrated, and scaled with XDS (54), leaving 5% of the reflections apart for cross-validation. The P9-1ΔC-arm structure was solved by molecular replacement with Phaser (55) using the coordinates of RBSDV P9-1 as a search model (PDB code: 3VJJ) (31). Refinement and manual model building were then performed with the programs Buster (56) and Coot (57), respectively. Due to the low resolution, on the initial refinement cycles specific reference-model restraints using RBSDV P9-1 as a template along with an automatic setting of the relative weight between geometry and X-ray terms were applied to assure the correctness of the model. Intermediate refinement steps performed on the PDB_REDO server (58) were critical for structure model optimizations. The final model was validated with MolProbity (36) as well as with the validation module implemented in Coot (57). **Table S2** summarizes the statistics generated at these steps.

### Prediction of IDRs

IDRs were predicted based on the P9-1 amino acid sequence (Uniprot D9U542) using a combination of the following servers: IUPred/Anchor (59), PONDR (VLXT and VSL2 mode) (http://www.pondr.com/) and PrDOS (60).

### Cryo-EM data acquisition of full-length P9-1, image processing, single-particle reconstruction, and refinement

High-quality recombinant P9-1 protein samples were suspended in buffer 10 mM Tris-HCl, 25 mM sodium chloride, pH 7.6, at 17 mg/ml and kept on ice before cryo-grid preparation. Several serial dilutions were prepared and 3 µl of each sample was loaded on Quantifoil R2/2Cu/Rh 300 holey-carbon-supported grids (Quantifoil Micro Tools GmbH, Jena, Germany). Initial cryo-EM sample preparations showed clear protein aggregation, which was reverted by omitting the glow-discharge step on the grids. The samples were incubated with the grids for 1 min, blotted by filter papers, and then plunge-frozen into liquid ethane cooled by liquid nitrogen using a Leica EM CPC manual plunger. The vitrified grids were stored in liquid nitrogen for later use. The best grids were obtained at 1/10 (v/v), where homogenous and well-spread individual particles were clearly identified. Data acquisition was performed using a Talos Arctica microscope (Thermo Fisher) operated at 200 kV with an FEI Falcon III direct detector at Centro Nacional de Biotecnología (CNB, Spain) for one day per grid using a nominal magnification of 120,000, corresponding to a calibrated pixel size of 0.855 Å per pixel and a defocus range of -0.8 to -3.8 μm. A total number of 669 micrograph movies were recorded under low dose conditions and fractionated into 60 frames each with a dose of 0.5 e-/Å^2^ per frame. All data processing was executed using Scipion (61), a software framework integrating several 3DEM software packages, as detailed below.

Micrographs were aligned for motion correction purposes and dose-weighted with MotionCor2 (62). Determination of the CTF, beam-induced movement, defocus values, astigmatism, and micrograph resolution estimation were performed using Ctffind4 (63). The final images were carefully examined for further image processing considering the particle distribution, the resolution, and the quality of Thon ring fitting. An initial template-free particle picking was performed (firstly manually and then automatically) using Xmipp (64). The preliminary set of picked single particles (202,824 particles) was first subjected to an initial 2D classification resulting in 163,663 particles with 5-fold symmetry and 30,526 with 6-fold symmetry (**Fig. S4**). Next, single 2D class averages were used as references using Eman2 (65) for getting two preliminary *ab-initio* volumes with D5 and D6 symmetries which were used as a reference for 3D classification and refinement. After several rounds of refinement two clearly different and well-populated 3D classes, decamer (with imposed D5 symmetry) and dodecamer (with imposed D6 symmetry) were found. The particles were then further extensively 3D classified using Relion-3 (66) resulting in a major population of 99,682 (decamer) and 22,510 (dodecamer) particles. Reconstructions of the final maps were sharpened by dividing the maps by the modulation transfer function of the detector and by applying a negative B-factor using Relion-3 (66). Local resolutions of the maps were calculated using ResMap (67). The data processing workflow is described in **Fig. S4**, and the data collection and reconstruction statistics are shown in **Table S3**.

### Molecular modeling, real-space refinement, and validation

The dimeric P9-1ΔC-arm crystal structure was fitted as a rigid body into the respective EM-density maps using UCSF ChimeraX (68). Later, the C-arm region (residues 314-337) was traced using the RBSDV P9-1 crystal structure (PDB code: 3VJJ) as a reference and manually adjusted using Coot (57). The docked atomic coordinates of the respective 3D models were refined into the locally filtered maps using phenix.real_space_refine with secondary structure restraints calculated in Phenix 1.18.2_3874 (69). The validation of the models was carried out using the MolProbity software (36). Model building and refinement statistics are shown in **Table S3**.

### SAXS analysis

P9-1 SAXS measurements were performed at the DO1B-SAXS1 beamline of the Brazilian Synchrotron Light Laboratory (LNLS, Brazil) with an incidence wavelength (λ) of 1.54 Å. The scattering intensity distributions as a function of the momentum transfer q were obtained in the q range between 0.013 and 0.48 Å^-1^ with q = 2π sin(θ)/λ where 2θ is the scattering angle. The SAXS patterns were recorded with exposure times of 20 s per frame for 10 min. A Pilatus 300K detector was used with an 883 mm sample detector distance. One-dimensional curves were obtained by integration of the 2D data using the program FIT-2D (70). Liquid samples were injected into the beamline vacuum-tight temperature-controlled X-ray cell for liquids. The P9-1 fractions obtained from exclusion chromatography were diluted from 5 to 0.5 mg/ml and no change in SAXS patterns with dilution was observed within this range of concentrations. Simulated patterns of the individual protein oligomers were done using pseudo-atom approximation obtained from low-resolution refinement of cryo-EM experiments with the combination of Scipion platform (61, 71) and ATSAS package (72). Simulated patterns were used for data interpretation employing least square procedures. Since a small proportion of larger aggregates were observed after purification, a fractal aggregate was included (73). Also, a Gaussian chain form factor (73) was used as a background function to account for flexible parts of the proteins. The OLIGOMER package (41) was also tested to estimate each oligomer volume fraction. *Ab initio* modeling was performed with the DENSS package (74) with imposed P2 symmetry.

### EMSA

Various amounts of purified P9-1, P9-1ΔC-arm, or BSA (as negative control) were incubated with 5 µM of a 22 nt HEX-DNA oligonucleotide probe (5’HEX-GACCTCGCTCTCTGTTTCTCAT 3’) in buffer 10 mM Tris-HCl, 50 mM potassium chloride, 0.5 mM ethylenediaminetetraacetic acid (EDTA), 10% glycerol, 1 mM dithiothreitol (DTT), pH 7.5 (20). Different concentrations of polyadenylic acid (polyA) (Midland Certified Reagent Company, USA) were employed for ssRNA competition experiments. According to the supplier, it consisted mostly of poly(A) polymers of more than 250 nucleotides in length. Reactions were held for 30 min at room temperature in a total volume of 20 µl and then subjected to 6.5% native PAGE. The migration of the labeled probe was detected in a XX6 G-box Imaging system (Syngene, USA). Three independent experiments were performed (n=3). The fluorescence intensity of the ssDNA probe in complex with the proteins was quantified with the ImageJ software (75) and the statistical significance of the signal was calculated using Two-way ANOVA followed by Tukey’s multiple comparisons test with GraphPad Prism 8 software (76).

### ATPase activity measurements

The initial rate of ATP hydrolysis for P9-1 and P9-1ΔC-arm was obtained from the slope of the time course of inorganic phosphate release. Reactions were carried out at 25 ºC in 25 mM Tris-HCl, 100 mM sodium chloride, 0.5 mM EDTA, 4.4 mM magnesium chloride, pH 7.7. All reactions were initiated with the addition of 2.5 mM ATP after protein preincubation for 10 min at 25 ºC in reaction media and a protein concentration of 6.5 µM. All reactions were stopped by the addition of ammonium heptamolybdate solution in an acidic medium and the amount of inorganic phosphate was quantified spectrophotometrically according to the Baginsky method (77) with modifications (78). The absorbance was measured in a Jasco V-550 spectrophotometer. When present in the reaction media, RNA was added before the 10 min protein preincubation. Five different reaction times were used to estimate the velocity and to ensure initial rate conditions the hydrolysis never exceeded 5% of the ATP present. The spontaneous hydrolysis of ATP was followed under identical conditions without protein and was negligible in these conditions.

### *In silico* reconstructions of the missing structural regions and MD simulations

The amino acid sequence of the full P9-1 dimer (674 residues) was aligned to the P9-1ΔC-arm crystal structure and a 3D model was created with MODELLER (79) to rebuild the regions not defined in the electron density map, and the C-arm residues (314 to 337). Protonation states at Ph 7.5 were assigned by the PDB2PQR server (80). The Cartesian coordinates of all residues present in the crystal structure were fixed in space as found experimentally by means of a harmonic potential of 10 kcal/mol/Å^2^; the remaining residues were allowed to move freely during all the following MD steps (see **Fig. 8A, *left*** and **Movie S1**). The model was minimized in implicit solvent, neutralized with 16 Na^+^ ions, solvated with explicit waters and 0.15 M NaCl, and minimized in solution. Then, it was thermalized to 299 K at NVT and simulated during 4 μs employing MD at NPT (P = 1 bar). The minimized model of the full-length dimer (D1) was aligned to each of the dimers in decamer (D5) and dodecamer (D6) to reconstruct the whole structures; residues from the C-arm region were replaced by those experimentally determined. The Cartesian coordinates of all residues experimentally defined were fixed in space using a harmonic potential of 10 kcal/mol/Å^2^; the remaining residues were allowed to move freely during all the following MD steps (see **Fig. 8A, *middle and right*** and **Movies S2** and **S3**). After minimization of the model *in vacuo*, some residues at the core of the structures and the C-arms were manually shifted to deinterlace regions from neighboring dimers, and a new minimization was run, removing the restraint over those residues. The models were then neutralized with 80 Na^+^ ions (decamer) and 96 Na^+^ ions (dodecamer), solvated with explicit waters and 0.15 M NaCl, and minimized in solution. Next, the systems were thermalized to 299 K at NVT and simulated during 3 μs (decamer) or 2 μs (dodecamer) by means of MD at NPT (P = 1 bar).

To treat every protein we used the ff19SB force field (81), and the entire system was surrounded by a truncated octahedral box of TIP3P water molecules (82), applying Dang’s parameters on ions (83). All systems were simulated using the Langevin algorithm to control the temperature and the pressure, with a coupling constant of 5 ps. SHAKE was used to keep all bonds involving hydrogen at their equilibrium values, which allowed us to use a 2 fs step for the integration of Newton’s equations of motion. Long-range electrostatic interactions were accounted for by using the Particle Mesh Ewald method with standard defaults. All simulations were carried out using the PMEMD CUDA code module of AMBER18 and analyzed with CPPTRAJ (84).

### Molecular Interaction Potentials

The linear PBE (without considering dielectric self-interaction), as implemented in CMIP (85), was used to compute free molecular interaction potentials using phosphate groups as probes. Experimentally-determined structures rebuilt and simulated by means of MD simulations were used as initial structures for the protein complexes. Representative structures were chosen from the two most populated clusters (based on the r.m.s.d fluctuations, after convergence), *i*.*e*. the two most prevalent conformations observed during MD simulations. The ionic strength was set to 0.15 M, and the reaction-field dielectric constants for proteins and water were set to 4.0 (86) and 79.8, respectively. The van der Waals radii were taken from the ff99SB force field (81).

### Graphics and molecular analyses

Graphs were plotted with GraphPad Prism 8 software (76). Structural analyses and figures were performed using PyMOL Molecular Graphics System 1.8 (Schrödinger, USA), UCSF ChimeraX (68), and VMD 1.9.3 (87). Movies were generated with Molywood (88).

## ACKNOWLEDGMENTS

This work was supported by the Argentinean Ministry of Science and Technology (MINCyT), the Argentinean Research Council (CONICET), and the National Agency for the Promotion of Science and Technology of Argentina (ANPCyT), under grants PICT 2014-3754, PICT 2015-0621, PICT 2016-1425, PICT 2017-2537 and PICT 2017-3150. We acknowledge the support of the European Union (EU) and Horizon 2020 through the iNEXT-Discovery Proposal 871037 (access project PID: 5391), the support and the use of resources of Instruct, a Landmark ESFRI project (specifically Instruct Access Project PID: 5526) and CRIOMECORR project (ESFRI-2019-01-CSIC-16) to the cryo-EM facility of the CNB (Spain). We would like to thank Prof. Dr. J. Dubcovsky (UC-Davis, USA) for kindly providing the pUC57-43 vector. We are also grateful to all members of the Nanotechnology National Laboratory (Brazil) for their additional help on cryo-EM data collection. CHI, MLC, SK, VA, FAG, MdV, and LHO are researchers from CONICET, while GL and MdV are researchers from INTA. DM and GS would like to acknowledge fellowship support from CONICET and ANPCyT, respectively. LHO acknowledges CONICET for travel support to cryo-EM facilities for data acquisition. We are grateful for the access to the PROXIMA-2A beamline at the SOLEIL Synchrotron (France) under the project 20170984. SAXS experiments were performed at the Brazilian Synchrotron Light Laboratory (Brazil) through the project SAXS1 – 20180371. AT and PDD acknowledge the graduate scholarship from CAP (UdelaR, Uruguay). PDD is an SNI (ANII) and PEDECIBA Química (UdelaR, Uruguay) researcher.

## Author Contributions

MdV and LHO designed and supervised the project. GL and DM designed the mutagenesis experiments. GL, VA and SMA performed the *in vivo* experiments. GL, DM and GS purified the proteins. GL and YGS carried out the analytical gel filtration experiments. GS and MLC executed the SEC-SLS measurements. CH-I performed and analyzed SAXS experiments, while DM, YGS and LHO assisted during the measurements and the analysis. MLC performed DLS measurements. GS and SK crystallized the protein, LHO performed the crystallographic data collection, processed the data, solved and refined the crystallographic structure. RA and LHO prepared the cryo-EM grids and performed the data collection. RM processed the cryo-EM data. LHO determined the 3D reconstructions and built and refined the atomic models. MdV and GL performed the nucleic acid binding assays. EM and SBK performed and analyzed the ATPase activity experiments. AT and PDD performed the theoretical modeling and MD simulations. GL, JMC, FAG, MdV and LHO financed the project. GL, DM, MdV and LHO analyzed, interpreted and discussed all results. GL, MdV and LHO wrote the paper with contributions from all authors.

## Accession numbers

P9-1ΔC-arm coordinates and structure factors were deposited in the Protein Data Bank (http://www.wwpdb.org/) with accession code 6UCT. Full-length MRCV P9-1 cryo-EM maps were deposited in the Electron Microscopy Data Bank (http://www.ebi.ac.uk/pdbe/emdb/) under the accession codes EMD-23046 (decamer D5) and EMD-23047 (dodecamer D6). The associated atomic models were deposited into the Protein Data Bank with accession codes 7KVC and 7KVD, respectively.

