## Supplementary Material for "A fijivirus major viroplasm protein shows RNA-stimulated ATPase activity by adopting pentameric and hexameric assemblies of dimers"

**This PDF file includes:**

Figs. S1 to S6
Tables S1 to S3

**Other Supplementary Materials for this manuscript include the following:**

Movies S1 to S3

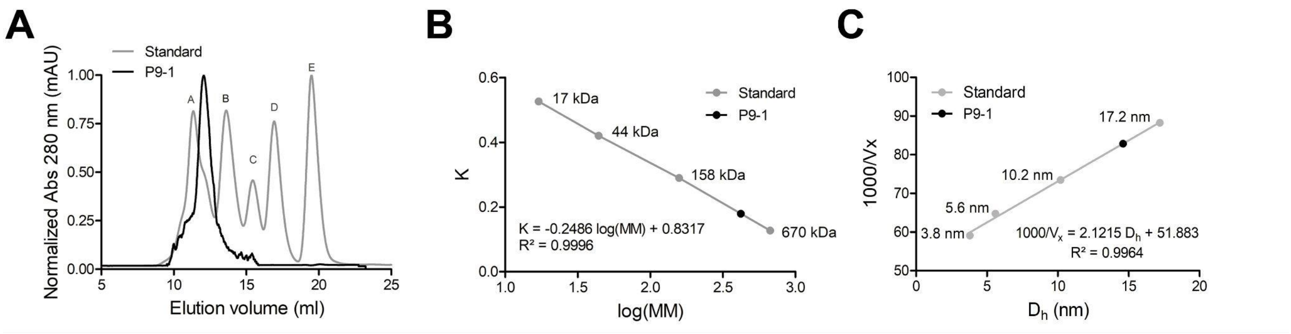


**Figure S1. Analytical SEC analysis of full-length P9-1.**

**A.** Chromatogram curves are shown for P9-1 (black) and for the standard protein samples (grey). Standard protein elution peaks are indicated as follows: peak A, bovine thyroglobulin MM = 670 kDa, $D_h$ = 17.2 nm; peak B, bovine γ-globulin, MM = 158 kDa, $D_h$ = 10.2 nm; peak C, chicken ovalbumin, MM = 44 kDa, $D_h$ = 5.6 nm; peak D, horse myoglobin, MM = 17 kDa, $D_h$ = 3.8 nm; and peak E, vitamin B12, MM = 1.35 kDa, $D_h$ = n/d. mAU: mili-UV Absorbance Units **B.** Estimation of the experimental MM of P9-1 (421.1 kDa) based on the partition coefficient (K). **C.** Estimation of the $D_h$ of P9-1 (14.6 nm) based on the elution volumes ($V_x$) and the $D_h$ of standard proteins.

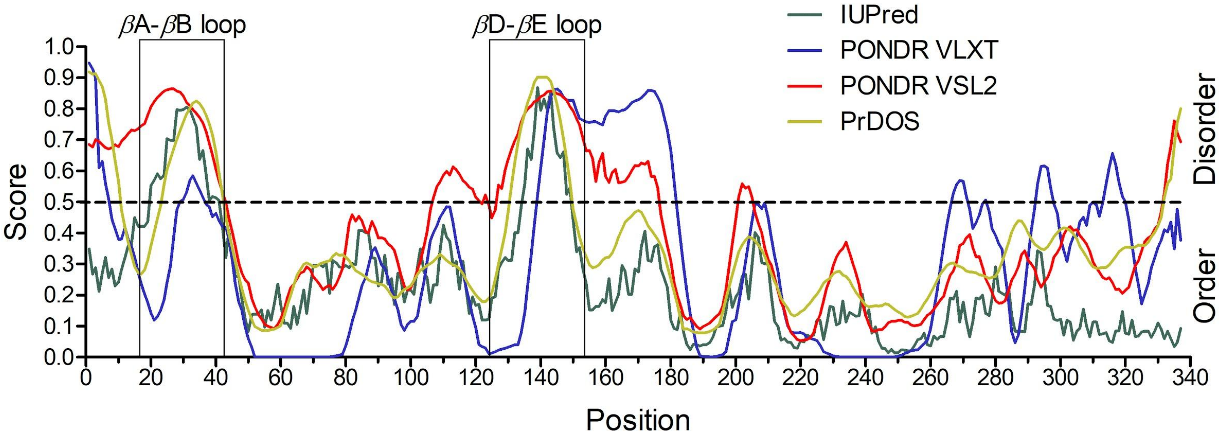


**Figure S2**. **Prediction of IDRs in the P9-1 sequence.**

Identification of IDRs according to the software IUPred2A (green line), PONDR VLXT (blue line), PONDR VLS2 (red line), and PrDOS (yellow line). Score values above the cut-off (0.5) indicate disordered residues. The regions comprising the loops βA-βB and βD-βE are highlighted in rectangles.

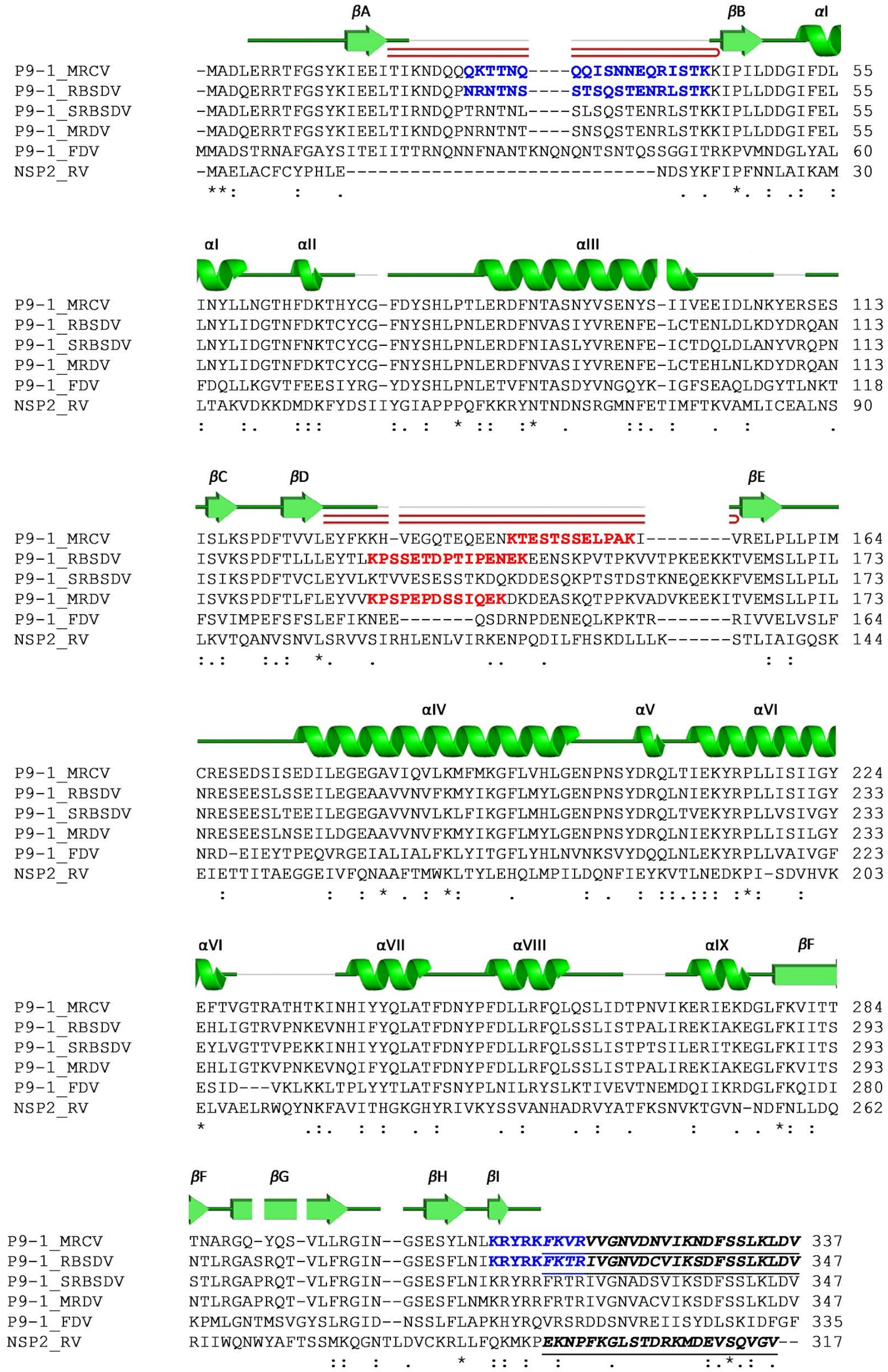


**Figure S3. Structure-sequence relationship of P9-1 and sequence alignment between fijivirus P9-1 proteins and rotavirus (RV) NSP2.**

Schematic diagrams of structural elements of P9-1 (PDB code: 6UCT) were obtained from PDBsum (89) depicted as spirals (α-helices I to IX), arrows (β-strands A to I), while loops are indicated as ⊃ in red. Multiple sequence alignment of P9-1 (UniProt D9U542, PDB code: 6UCT) with RBSDV (UniProt Q913E4, PDB code: 3VJJ), SRBSDV (UniProt B6SCH3, PDB code: 5EFT), MRDV (UniProt A0A650ABG4), FDV (UniProt Q9YX38) counterparts and RV NSP2 (UniProt Q03243, PDB code: 1L9V) was performed using Clustal Omega (90). Potential RNA binding residues are indicated in bold and blue, while PEST sequences are in bold and red. The C-arm of MRCV and RBSDV, and the NSP2 C-terminal region (CTR) are underlined in bold and italics.

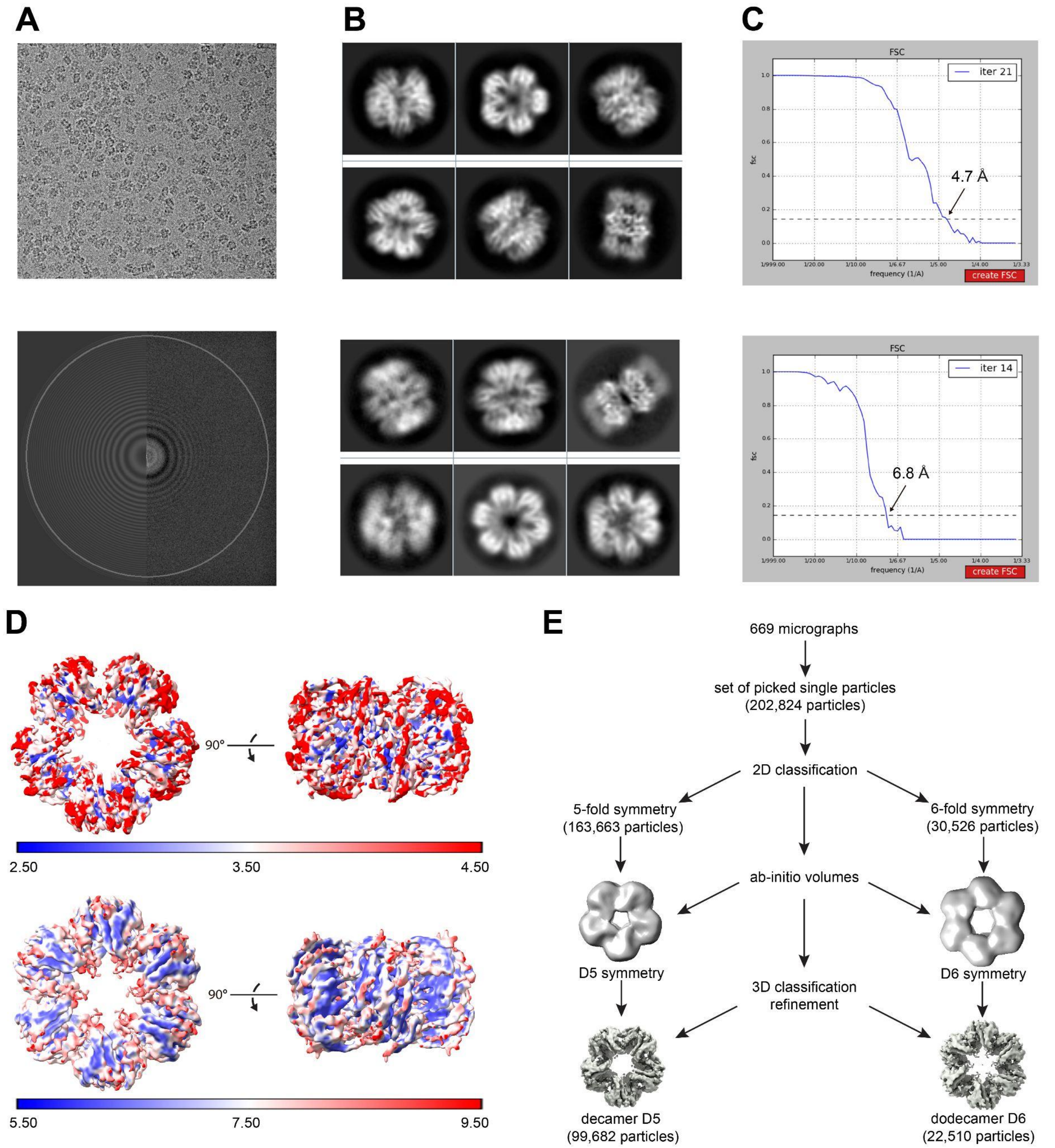


**Figure S4. Single-particle cryo-EM data processing and validation of full-length P9-1.**

**A.** Representative motion-corrected cryo-electron micrograph (*upper*). Fourier transformation showing visible thon rings (*bottom*). **B.** Reference-free 2D class average of decamer D5 symmetry (*upper*) and dodecamer D6 symmetry (*bottom*). **C.** Gold-standard Fourier shell correlation (FSC) curves for the decamer (*upper*) and dodecamer (*bottom*). The

0.143 cut-off is indicated by a horizontal dashed black line. **D.** Local resolution map for the decamer (*upper*) and dodecamer (*bottom*). **E.** Cryo-EM data processing flow-chart.

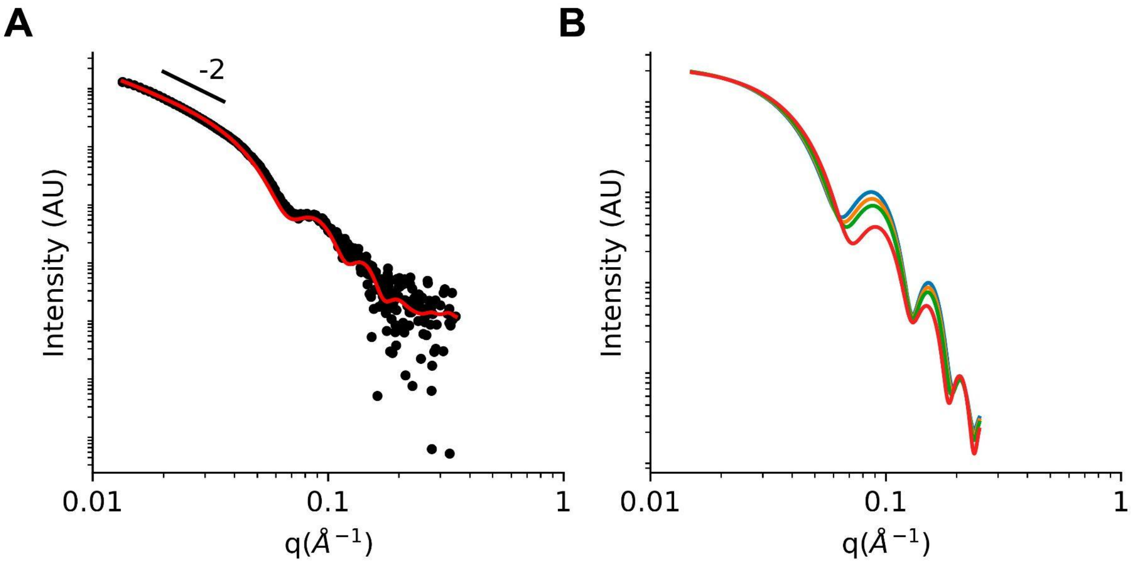


**Figure S5**. **SAXS patterns in function of the momentum transfer q.**

**A.** Log-log SAXS pattern from a largely aggregated fraction obtained from SEC. The slope at low angle was -2. Dots represent experimental data and the fitted curve is depicted in a continuous red line. AU, arbitrary units. **B.** Contrast effect in simulated corona form factor. Empty corona represents the case of equal density contrast between porous and solvent (blue) and in red line the fulfilled internal pore. This simulation shows the filling effects of the internal pore over the form factor where water and ions or small organic molecules could affect harmonics positions and their relative intensities.

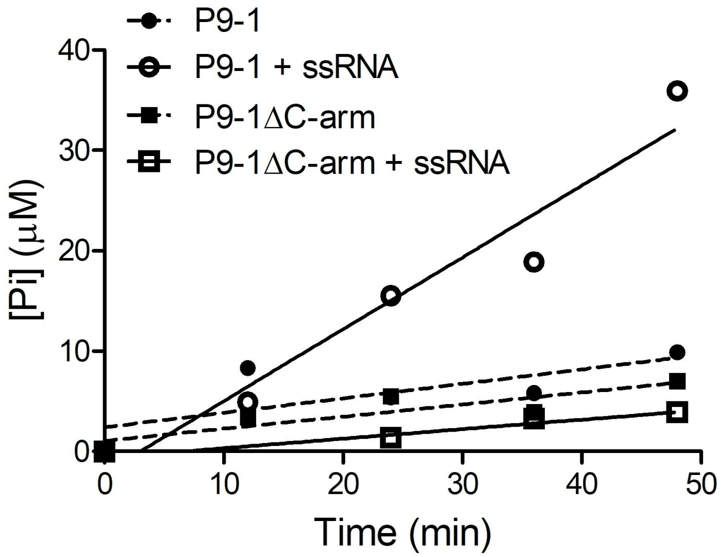


**Figure S6. Time courses of the release of inorganic phosphate from ATP catalyzed by P9-1 and P9-1ΔC-arm and the effect of ssRNA.**

Continuous lines are graphical representations of linear functions fitted to the experimental data by linear regression analysis. Experiments were carried out at 25 ºC and contained 6.5 μM protein, 2.5 mM ATP, and either 0 or 500 μM ssRNA, in reaction media consisting of 25 mM Tris-HCl, 100 mM sodium chloride, 0.5 mM EDTA, 4.4 mM magnesium chloride, pH 7.7.

**Table S1. DLS size distribution analysis of P9-1 and P9-1ΔC-arm proteins.**

| *Sample* | *Z-Average (nm)* | *PdI* | *$D_h$ Num (nm)* | *% Mass* |
|---|---|---|---|---|
| P9-1 | 27.98 ± 1.47 | 0.435 ± 0.015 | 14.17 ± 0.87 | 99.7 ± 0.2 |
| P9-1ΔC-arm | 18.24 ± 0.27 | 0.333 ± 0.014 | 8.41 ± 1.40 | 99.7 ± 0.5 |

PdI: polydispersity index. $D_h$ Num: diameter in the number distribution. % Mass: % area in the volume distribution.

**Table S2. X-ray diffraction data collection and refinement statistics.**

| | |
|---|---|
| *Data collection* | |
| Wavelength (Å) | 0.9801 |
| Crystal-detector distance (mm) | 241.88 |
| Rotation range per image (°) | 0.1 |
| No. of frames | 3600 |
| Exposure time per image (s) | 0.025 |
| *Indexing and scaling* | |
| Cell parameters | |
| a=b (Å) | 86.56 |
| c (Å) | 95.60 |
| α=β=γ (°) | 90 |
| Space group | $P4_32_12$ |
| Mosaicity (°) | 0.17 |
| Resolution range (Å) | 47.80 – 3.47 |
| Total No. of reflections | 125433 (29122) |
| No. of unique reflections | 5035 (1138) |
| Completeness (%) [a] | 99.2 (96.8) |
| Redundancy | 24.9 (25.6) |
| ⟨I/σ(I)⟩ | 15.8 (1.6) |
| $R_{meas}$ | 0.137 (3.019) |
| $R_{pim}$ | 0.027 (0.582) |
| $CC_{1/2}$ (%) | 0.999 (0.548) |
| Solvent content (%) | 49 |
| No. of chains per asymmetric unit | 1 |
| Overall *B* factor from Wilson plot ($Å^2$) | 111 |
| *Refinement* | |

| | |
|---|---|
| Resolution range (Å) | 43.28 – 3.47 |
| Number of protein atoms | 2013 |
| Number of ligand atoms | - |
| Number of water molecules | - |
| *R* | 0.220 |
| $R_{free}$ | 0.277 |
| R.m.s. deviations from ideal values (91) | |
| Bond lengths (Å) | 0.010 |
| Bond angles (º) | 1.04 |
| Average *B* factor ($Å^2$) | 185 |
| *Validation* (36) | |
| MolProbity score (percentile) | 2.53 ($98^{th}$*) |
| Ramachandran plot | |
| Favored (%) | 91.3 |
| Allowed (%) | 7.4 |
| Disallowed (%) | 1.3 |
| Cβ outliers (%) | 0 |
| CaBLAM outliers (%) | 1.4 |
| Cα geometry outliers (%) | 0 |
| PDB code | 6UCT |

[a] Values for the outer shell are given in parentheses (3.80 – 3.47 Å).
[*] $100^{th}$ percentile indicates the best structures of comparable resolution; $0^{th}$ percentile indicates the worst ones.

**Table S3. Cryo-EM data collection, refinement and validation statistics.**

| | *P9-1 decamer (D5)* | *P9-1 dodecamer (D6)* |
|---|---|---|
| *Data collection and processing* | | |
| Microscope | Talos Arctica | Talos Arctica |
| Voltage (kV) | 200 | 200 |
| Detector | FEI Falcon III | FEI Falcon III |
| Magnification | 120,000 | 120,000 |
| Electron exposure ($e^{-}/Å^2$) | 30 | 30 |
| Defocus range (μm) | −0.8 to −3.8 | −0.8 to −3.8 |
| Pixel size (Å) | 0.855 | 0.855 |
| Symmetry imposed | *D*5 | *D*6 |
| Initial particle images (no.) | 202,824 | 202,824 |
| Final particle images (no.) | 99,682 | 22,510 |
| Map resolution (Å) | 4.7 | 6.8 |
| FSC threshold | 0.143 | 0.143 |
| EMD code | EMD-23046 | EMD-23047 |
| *Model Refinement* | | |
| Initial model used (PDB code) | 6UCT | 6UCT |
| Unmasked resolution at 0.5/0.143 FSC (Å) | 5.3/4.7 | 7.1/5.6 |
| Masked resolution at 0.5/0.143 FSC (Å) | 5.2/4.6 | 7.0/5.6 |
| Number of protein atoms | 21,920 | 26,304 |
| Number of ligand atoms | - | - |
| Number of water molecules | - | - |
| *B* factors ($Å^2$) | 170 | 175 |
| R.m.s. deviations from ideal values (91) | | |
| Bond lengths (Å) | 0.005 | 0.004 |
| Bond angles (°) | 0.846 | 1.008 |

| | | |
|---|---|---|
| *Validation (36)* | | |
| MolProbity score (percentile) | 2.57 (43$^{rd}$*) | 2.52 (46$^{th}$*) |
| CC (mask) | 0.82 | 0.75 |
| Ramachandran plot | | |
| Favored (%) | 89.3 | 91.2 |
| Allowed (%) | 10.7 | 8.8 |
| Disallowed (%) | 0 | 0 |
| Cβ outliers (%) | 0 | 0 |
| CaBLAM outliers (%) | 1.7 | 2.5 |
| Cα geometry outliers (%) | 0.4 | 0.4 |
| PDB code | 7KVC | 7KVD |

* 100$^{th}$ percentile indicates the best structures of comparable resolution; 0$^{th}$ percentile indicates the worst ones.

***Movie_S1. (separate file)***

**Animation of the reconstructed model of the P9-1 dimer and its MD simulation.** The model is colored according to secondary structure elements as in **Fig. 3A**. When the MD animation starts, residues that were fixed in space as found experimentally are colored in gray. The MD animation displays the last 3 μs of a 4 μs simulation.

***Movie_S2. (separate file)***

**Animation of the reconstructed model of the P9-1 decamer and its MD simulation.** The model is colored according to its secondary structure elements as in **Fig. 3A**. At the beginning of the movie, only one dimer is shown and the other four appear sequentially to illustrate the decamer construction. Then, the MD animation starts, and residues that were fixed in space as found experimentally are colored in gray using Connolly surface. The structure is zoomed-in to highlight the mobility at the inner pore of the complex. Then, a parallel plane to the pore is presented (three out of the five dimers are shown in this last segment). The MD animation displays the last 2 μs of a 3 μs simulation.

***Movie_S3. (separate file)***

**Animation of the reconstructed model of the P9-1 dodecamer and its MD simulation.** The model is colored according to its secondary structure elements as in **Fig. 3A**. At the beginning of the movie, only one dimer is shown, and the other five appear sequentially to illustrate the dodecamer construction. Then, the MD animation starts, and residues that were fixed in space as found experimentally are colored in gray using Connolly surface. The structure is zoomed-in to highlight the mobility at the inner pore of the complex. Then, a parallel plane to the pore is presented (three out of the six dimers are shown in this last segment). The MD animation displays a 2 μs simulation.
